# Endophilin A1 promotes Actin Polymerization in response to Ca^2+^/calmodulin to Initiate Structural Plasticity of Dendritic Spines

**DOI:** 10.1101/2020.05.07.082081

**Authors:** Yanrui Yang, Jiang Chen, Xue Chen, Di Li, Jianfeng He, Shun Zhao, Xiaoyu Yang, Shikung Deng, Dou Wang, Zhenzhen Guo, Shaoxia Zhu, Dong Li, Cong Ma, Xin Liang, Yun S. Shi, Jia-Jia Liu

## Abstract

Dendritic spines of excitatory neurons undergo activity-dependent structural and functional plasticity, which are cellular correlates of learning and memory. However, mechanisms underlying the rapid morphological changes immediately after NMDAR-mediated Ca^2+^ influx into spines remain poorly understood. Here we report that endophilin A1, a neuronal N-BAR protein, orchestrates membrane dynamics with actin polymerization to initiate spine enlargement in the induction phase of long-term potentiation (LTP). Upon LTP induction, Ca^2+^/calmodulin enhances its binding to both membrane and p140Cap, a cytoskeleton regulator. As a result, endophilin A1 rapidly associates with the relaxed plasma membrane and promotes actin polymerization, leading to acute expansion of spine head. Moreover, not only the p140Cap-binding, but also calmodulin- and membrane-binding capacities of endophilin A1 are required for LTP and long-term memory. Thus, endophilin A1 functions as calmodulin effector to drive spine enlargement in response to Ca^2+^ influx in the initial phase of structural plasticity.

## Introduction

Long-term potentiation (LTP) of synaptic strength contributes to neural mechanisms underlying learning and memory (Nabavi et al., 2014). In the mammalian brain, most glutamatergic synapses are located on dendritic spines, micron-sized protrusions from dendrites. In response to input activity, spines undergo changes in both morphology (structural plasticity) and function (functional plasticity), which are tightly correlated during LTP (Harvey and Svoboda, 2007; Matsuzaki et al., 2001; Matsuzaki et al., 2004). Imaging studies have revealed that induction of LTP triggers a large transient increase in spine volume (1-5 min after stimulation, early or transient phase) that decays to a long-lasting spine size expansion (> 40 min, late or sustained phase) (Harvey and Svoboda, 2007; Matsuzaki et al., 2004), a process termed structural LTP (sLTP), which would allow physical enlargement of glutamatergic synapses to accommodate more α-amino-3-hydroxy-5-methyl-4-isoxazolepropionic acid receptors (AMPARs) for synaptic potentiation (Herring and Nicoll, 2016). Moreover, recent studies have established a direct link between spine morphological changes and memory trace *in vivo* by demonstrating disruption of acquired motor learning by optical shrinkage of potentiated spines in the motor cortex (Hayashi-Takagi et al., 2015). Although sLTP has been studied intensively, with calcium signaling-regulated actin remodeling being the central process that governs the stabilization and consolidation of spine enlargement (Nakahata and Yasuda, 2018), mechanism(s) initiating rapid spine head expansion remains largely unexplored due to limited spatiotemporal resolution of the molecular events in the acute phase (0-1 min) of LTP induction.

Endophilin A1 is a member of the endophilin A protein family characterized by an N-terminal BIN/amphiphysin/Rvs (BAR) domain and a C-terminal Src homology 3 (SH3) domain. The gene encoding endophilin A1 (*EEN1*, a.k.a *sh3gl2*) is almost exclusively expressed in brain (Ringstad et al., 1997) and has been implicated in epilepsy, Alzheimer’s disease and schizophrenia (Corponi et al., 2019; Ren et al., 2008; Yu et al., 2018a; Yu et al., 2018b). Originally identified as a component of the endocytic machinery, endophilin As function in synaptic vesicle recycling at the presynaptic site in two distinct processes: ultrafast endocytosis from the plasma membrane following synaptic vesicle fusion and clathrin uncoating of regenerated synaptic vesicles (Milosevic et al., 2011; Ringstad et al., 1997; Schuske et al., 2003; Verstreken et al., 2003; Watanabe et al., 2018). Recently endophilin A2 has also been found to mediate fast clathrin-independent endocytosis in mammalian epithelial cells (Boucrot et al., 2015; Renard et al., 2015). Other studies have implicated endophilin As in autophagosome formation and protein homeostasis at presynaptic terminals (Murdoch et al., 2016; Soukup et al., 2016). In dendrites, both endophilin A2 and A3 interact with Arc/Arg3.1 to accelerate endocytosis of AMPARs at the postsynaptic membrane during late-phase synaptic plasticity (Chowdhury et al., 2006).

Previously we found that during synaptic development, endophilin A1 contributes to dendritic spine morphogenesis and stabilization through interaction with p140Cap, an actin cytoskeleton regulator (Yang et al., 2015). We also found that *EEN1* gene knockout (KO) in the hippocampal CA1 region of mouse brain causes impairment of LTP of the Schaffer collateral-CA1 pathway and long-term memory (Yang et al., 2018). At the cellular level, endophilin A1, not A2 or A3, is required for N-methyl-D-aspartate receptor (NMDAR)-mediated synaptic potentiation of dendritic spines in mature CA1 pyramidal cells (Yang et al., 2018). Intriguingly, overexpression of p140Cap, its downstream effector, fails to rescue the structural and functional plasticity of spines in *EEN1* KO (*EEN1*^*−/−*^) neurons (Yang et al., 2018), suggesting the necessity of spatiotemporal coordination of membrane expansion and actin polymerization during synaptic potentiation. In this study, we investigated the mechanistic role(s) of endophilin A1 in sLTP with a combination of cell biological, biochemical, electrophysiological and genetic approaches. We present evidence that endophilin A1 serves as an immediate effector of Ca^2+^/calmodulin to promote actin polymerization-dependent membrane expansion in the initial phase of spine structural plasticity.

## Results

### Endophilin A1 is Required for the Acute Structural Plasticity of Dendritic Spines

Knockout of the *EEN1* gene causes inhibition of structural and functional plasticity of dendritic spines in hippocampal neurons (Yang et al., 2018). To investigate the mechanistic role(s) of endophilin A1 in synaptic plasticity, first we determined at which temporal stage(s) of LTP it functions by rescuing the morphological phenotype of *EEN1* KO neurons with overexpressed endophilin A1 in the induction phase of chemically-induced LTP (cLTP) (Figure 1A). Quantification of spine size indicated that endophilin A1 is required for spine enlargement as early as 1 minute after application of the NMDAR co-agonist glycine (Figure 1B and 1C).

**Figure 1.**
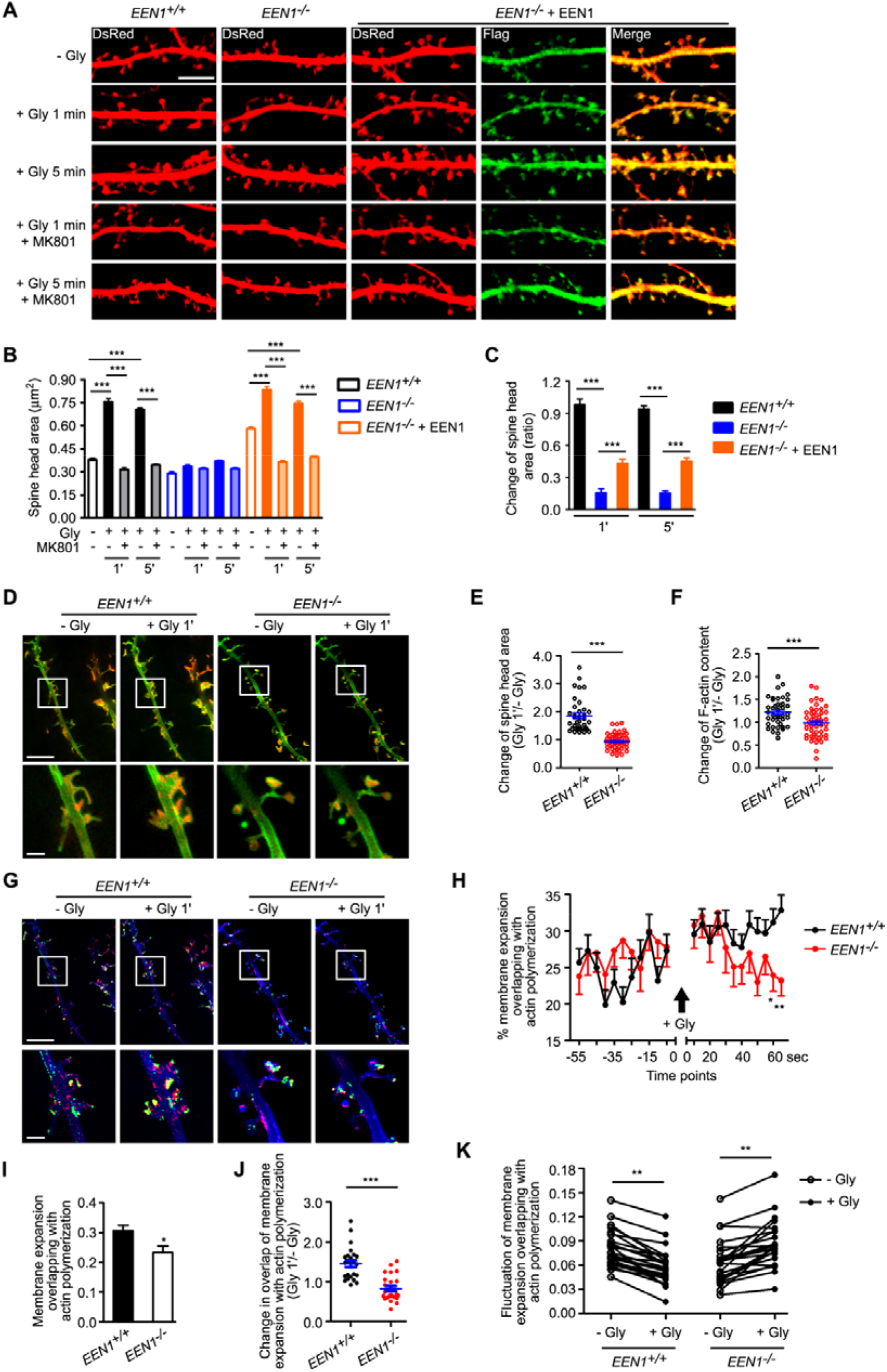
Endophilin A1 is required for spine expansion and actin polymerization in the initial Phase of sLTP. (**A**) Mouse hippocampal neurons were co-transfected with pLL3.7-DsRed (volume marker) and pCMV-Tag2B (Flag vector) or pCMV-Tag2B-endophilin A1 (Flag-EEN1) on DIV12, pre-treated with DMSO (vehicle control) or MK801 (NMDAR antagonist) and chemically induced LTP with glycine (Gly) on DIV16. Neurons were fixed 1 min or 5 min after glycine application, stained with antibodies to Flag and imaged by confocal microscopy. Bar = 5 μm. (**B**) Quantification of spine size in (**A**). N ≥ 13 cells, n ≥ 563 spines per group. (**C**) Changes of spine size in (**A**). (**D**) GI-SIM imaging of *EEN1* WT and KO hippocampal neurons expressing mGFP and LifeAct-mCherry. Glycine was applied after imaging for 1 min and imaging was continued for 5 more minutes. Shown are representative still images right before and 1 min after glycine application. Lower panels are magnification of boxed areas in upper panels. Bars: 5 μm (upper) and 1 μm (lower). (**E, F**) Quantitative analysis of spine size increase (**E**) or F-actin enrichment (**F**) in spines imaged by GI-SIM. (**G**) Spine membrane expansion overlapping with actin polymerization in D. Spine growth and increase in F-actin signals are color coded green and red, respectively. Bars: 5 μm (upper) and 1 μm (lower). (**H**) Quantification of fractions of expanded membrane overlapping with actin polymerization in individual spines of *EEN1* WT and KO neurons before and after glycine application at 5 s intervals. (**I**) Quantification of fractions of membrane expansion overlapping with actin polymerization at 1 min after glycine application in individual spines of *EEN1* WT and KO neurons. (**J**) Changes in the extent of overlap between membrane expansion and actin polymerization 1 min after glycine application in individual spines of *EEN1* WT and KO neurons. Data are normalized to the time point right before glycine application. (**K**) Mean fluctuation of the overlap between membrane expansion and actin polymerization in individual spines within 1 min before and after glycine application for *EEN1* WT and KO neurons. Data represent mean ± SEM in (**B**), (**C**) and (**I**). WT: N = 5 neurons and n = 24 spines; KO: N = 6 neurons and n = 27 spines in (**E**), (**F**) and (**H-K**). * *p* < 0.05, ** *p* < 0.01, *** *p* < 0.001.

Actin polymerization in dendritic spines is crucial for not only spine morphogenesis during synaptic development but also sLTP of mature neurons. Imaging studies on hippocampal neurons have shown that actin polymerization in spines starts as early as 20 s after LTP induction (Okamoto et al., 2004). As endophilin A1 is required for acute LTP-induced spine enlargement, we reasoned that it might function to promote actin polymerization in the initial phase of structural plasticity. To monitor morphological changes and actin dynamics of spines simultaneously, we performed super-resolution live imaging of *EEN1* wild-type (WT) and KO neurons expressing membrane-anchored GFP (mGFP) and the F-actin probe LifeAct-mCherry by Grazing Incidence Structured Illumination Microscopy (GI-SIM) (Guo et al., 2018). In WT neurons, consistent with our previous study (Guo et al., 2018), we observed rapid increase in both spine size and F-actin signal intensity in dendritic spines within 1 minute upon glycine application (Video 1 and Figure 1D-1F). In contrast, no significant changes in spine size were detected in *EEN1* KO neurons even though the shape of spines changed constantly (Video 2, Figure 1D and 1E). These live imaging data indicate that endophilin A1 is required for spine head enlargement during the initial phase of sLTP. Notably, although the spine heads of *EEN1* KO neurons were as motile as those of WT cells, the LTP-induced net increase in F-actin content was also abolished (Figure 1D and 1F), indicating that endophilin A1 is also required for spine actin polymerization in the initial phase of sLTP.

Consistent with previous findings (Guo et al., 2018; Honkura et al., 2008), we observed membrane expansion of spine head accompanied by local increase in F-actin content in the initial phase of sLTP (Figure 1G and Video 1). Quantitative analysis of the images clearly revealed that actin polymerization and plasma membrane protrusion of spine head are tightly coupled spatially and temporally (Figure 1H-K and Video 3), suggesting that local actin polymerization provides propulsive force for spine growth. Notably, compared with WT, the membrane protrusion of spine head and increase in F-actin content was much less coupled in *EEN1* KO neurons (Figure 1H-1K and Video 4), implicating endophilin A1 in actin polymerization-dependent membrane expansion in the initial phase of sLTP.

### Membrane Unfolding and Branched Actin Polymerization are Required for Acute Structural Plasticity

The observation that endophilin A1 is required for rapid spine membrane expansion upon glycine application prompted us to investigate mechanism(s) underlying initiation of sLTP. Although it was postulated that the membrane source for LTP-induced spine enlargement comes from transport of intracellular recycling endosomes to the neuronal plasma membrane (Park et al., 2004), imaging studies revealed that spine head expansion precedes most of the AMPAR exocytotic events during the early phase of cLTP (Kopec et al., 2006), and that the light chain of botulinum toxin type B (BoTox), a neurotoxin that binds to the SNARE complex and inhibits exocytosis, had no effect on the initial spine expansion after the theta burst paring protocol of LTP induction (Yang et al., 2008). To determine whether spine membrane expansion requires fusion of exocytosed vesicles with the plasma membrane in cLTP, we tested the effect of Tetanus toxin (TeTx), another inhibitor of SNARE-mediated membrane fusion, on glycine-induced acute increase in spine size. Consistent with previous studies (Hiester et al., 2018), treatment of hippocampal neurons with TeTx did not affect spine enlargement in the first 3 minutes of glycine treatment (Figure 2A and 2B), indicating that vesicle fusion is not the direct source of membrane supply for the rapid structural expansion of spines in the initial phase of sLTP.

**Figure 2.**
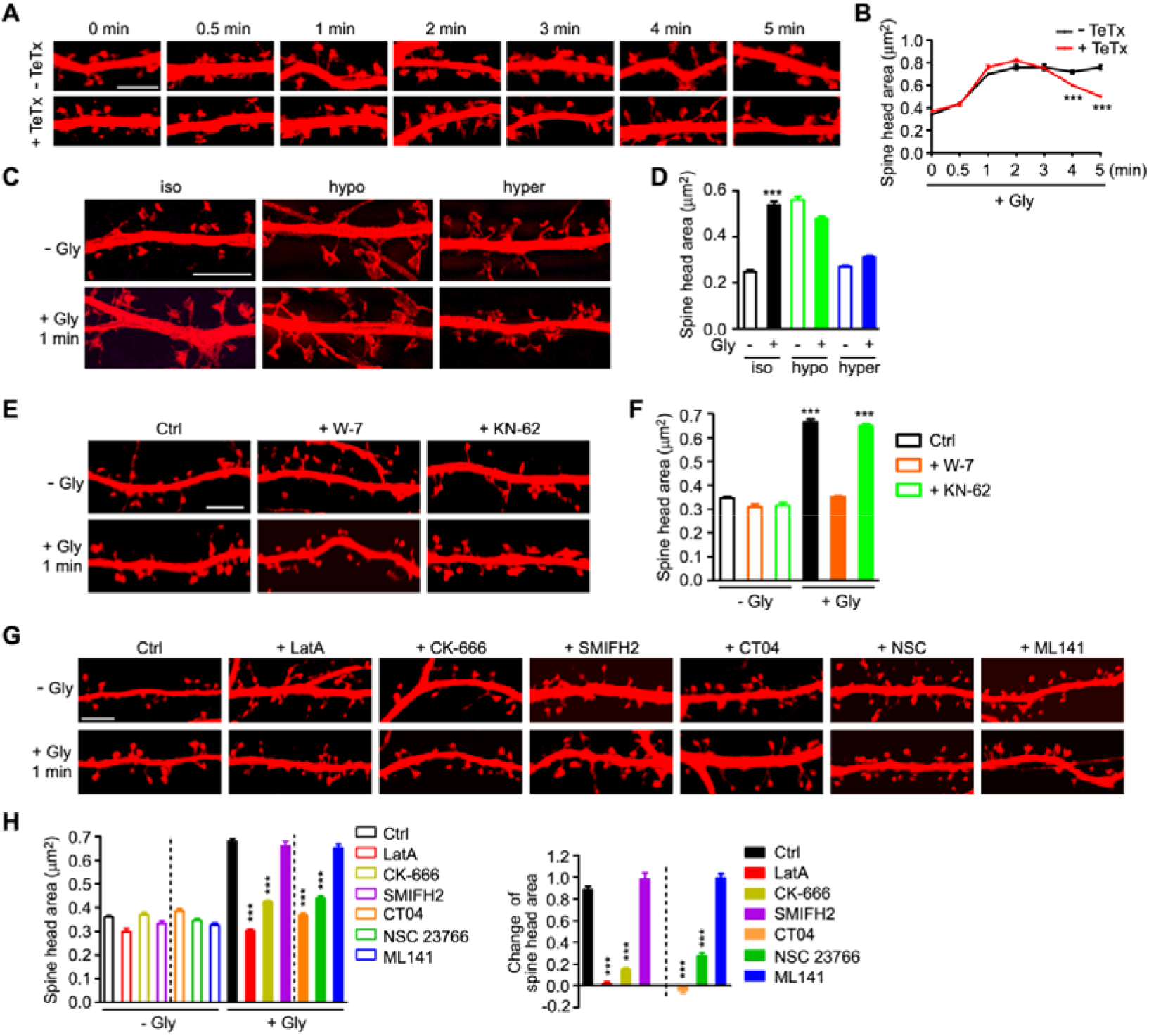
Membrane unfolding and branched actin polymerization are required for initiation of sLTP. (**A**) Mouse hippocampal neurons transfected with pLL3.7-DsRed on DIV12 were pre-treated with DMSO or tetanus toxin (TeTx) for 10 min and chemically induced LTP with glycine on DIV16. Neurons were fixed at different time points after glycine application and imaged by confocal microscopy. Bar = 5 μm. (**B**) Quantification of spine size in (**A**). N ≥ 12 neurons, n ≥ 530 spines per group, *** *p* < 0.001. (**C**) Effect of osmotic shock on spine enlargement 1 min after glycine application. Neurons were pre-treated with iso-osmotic, hypo-osmotic or hyper-osmotic solution respectively for 10 min and chemically induced LTP in the same solution on DIV16. Neurons were fixed 1 min after glycine application and imaged by 3D-SIM microscopy. Bar = 4 μm. (**D**) Quantification of spine size in (**C**). N ≥ 12 neurons, n ≥ 382 spines per group. (**E**) Effects of W-7 (calmodulin inhibitor) and KN-62 (CaMKII inhibitor) on spine enlargement 1 min after glycine application. Bar = 5 μm. (**F**) Quantification of spine size in (**E**). N ≥ 12 neurons, n ≥ 504 spines per group. (**G**) Effects of inhibitors for actin remodeling regulators on spine enlargement 1 min after glycine application. Shown are representative confocal images. Bar = 5 μm. CK-666: Arp2/3 inhibitor. SMIFH2: Formin inhibitor. CT04: RhoA inhibitor. NSC 23766: Rac1 inhibitor. ML 141: Cdc42 inhibitor. (**H, I**) Quantification of spine size (left) and changes in spine size (right) in (**G**). N ≥ 12 neurons, n ≥ 516 spines per group. Data represent mean ± SEM in (**B**), (**D**), (**F**) and (**H**). *** *p* < 0.001.

Both electron microscopy and super-resolution imaging reveal that the surface of mature spines is not smooth but rather convoluted (Arellano et al., 2007; Harris and Stevens, 1989; Smith et al., 2014). To determine whether the membrane folds or invaginations in spine head contribute to structural expansion, we increased membrane tension by exposing neurons to hypo-osmotic buffer and found that cLTP-induced spine enlargement was abolished (Figure 2C and 2D). As membrane tension increases fusion efficiency (Kliesch et al., 2017), these findings corroborate that membrane fusion does not contribute to rapid spine enlargement. Conversely, incubation of neurons with hyper-osmotic buffer, which shrunk the spines and generated membrane folds, also antagonized cLTP-induced spine enlargement (Figure 2C and 2D). These data together indicate that, similar to the formation of membrane expansion in migrating primordial germ cells (Goudarzi et al., 2017), the membrane supply for rapid spine enlargement in the initial phase of sLTP comes from plasma membrane invaginations, which are local unfolding of spine surface convolutions.

Previous studies on activity-dependent structural remodeling of dendritic spines indicate that glutamate uncaging-induced rapid spine enlargement requires NMDAR, calmodulin and actin polymerization, whereas long-lasting spine enlargement also requires the activity of Ca^2+^/calmodulin-dependent protein kinase II (CaMKII) (Matsuzaki et al., 2004). Indeed, inhibition of calmodulin but not CaMKII abolished glycine-induced spine enlargement of hippocampal neurons during the initial phase of sLTP (Figure 2E and 2F). The Ca^2+^/calmodulin◻CaMKII pathway triggers several signaling cascades to promote actin polymerization and AMPAR trafficking to the plasma membrane during synaptic potentiation (Murakoshi and Yasuda, 2012). In line with previous studies (Honkura et al., 2008; Matsuzaki et al., 2004), inhibition of actin polymerization with latrunculin A (LatA) abolished enlargement of spine head during the initial phase of sLTP (Figure 2G and 2H). Moreover, inhibition of Arp2/3 had a similar effect to that of LatA, whereas inhibition of Formin had no effect (Figure 2G and 2H), indicating that branched rather than linear actin polymerization is required for acute spine expansion.

The Rho family members of small GTPases are known downstream effectors of Ca^2+^/calmodulin and regulators of actin reorganization and structural plasticity (Hedrick and Yasuda, 2017; Spiering and Hodgson, 2011). Next we determined whether they are required for cLTP-induced acute spine enlargement by treating hippocampal neurons with inhibitor for RhoA, Rac1 or Cdc42. In agreement with previous findings that rapid structural remodeling requires actin severing and nucleation by ADF/cofilin, inhibition of RhoA, the upstream activator of ADF/cofilin (Hedrick et al., 2016; Murakoshi et al., 2011), abolished glycine-induced spine enlargement (Figure 2G and 2H). Notably, although both Rac1 and Cdc42 can activate actin polymerization (Hedrick and Yasuda, 2017), only Rac1 is required for the initial spine growth of sLTP (Hedrick et al., 2016; Murakoshi et al., 2011) (Figure 2G and 2H, this study). Moreover, inhibition of Rac1 only partially inhibited acute spine expansion (Figure 2G and 2H), suggesting the presence of other factor(s) that promotes branched actin polymerization in response to Ca^2+^ influx immediately upon sLTP induction. Collectively, these data indicate that Ca^2+^/calmodulin-regulated branched actin polymerization plays an essential role in acute spine membrane expansion in the initial phase of sLTP.

### Ca^2+^/calmodulin Enhances Endophilin A1-p140Cap Interaction to Promote Actin Polymerization in Spines

Having established that endophilin A1 is required for both spine head enlargement and actin polymerization in the initial phase of sLTP, we speculated that endophilin A1 promotes actin polymerization in spines to provide pushing force for membrane expansion. Since CaMKII is not required for acute spine enlargement, we reasoned that endophilin A1 might function earlier than CaMKII in molecular events triggered by NMDAR-mediated Ca^2+^ influx upon LTP induction. Previously studies reported that Ca^2+^ binding changes the conformation of endophilin A2 and regulates its interaction with dynamin and voltage-gated Ca^2+^ channels (VGCC) (Chen et al., 2003). Nevertheless, no direct binding between endophilin A1 and Ca^2+^ was detected by isothermal titration calorimetry (ITC) and fluorescence spectrometry (Figure 3-figure supplement 1).

Recent studies reported that calmodulin binds to mammalian N-BAR proteins including endophilin A1 and A2 (Myers et al., 2016). Indeed, GST-pull down assay showed that endophilin A1 binds to calmodulin via its N-terminal BAR domain and the interaction is strengthened by Ca^2+^ (Figure 3A-3F). We reasoned that Ca^2+^/calmodulin might function as upstream regulator of endophilin A1 function(s). As expected, co-immunoprecipitation (co-IP) from both transiently transfected HEK293 cells and mouse hippocampal neurons showed that the interaction between endophilin A1 and p140Cap is Ca^2+^-dependent (Figure 3G-3J). Moreover, co-IP from cultured hippocampal neurons revealed that LTP induction enhanced the interaction between endophilin A1 and p140Cap in a Ca^2+^- and NMDAR-dependent manner (Figure 3K and 3L). Further, the association of endophilin A1 with not only calmodulin but also p140Cap was enhanced acutely upon LTP induction, which was abolished with the calmodulin inhibitor W-7 (Figure 3M and 3N), indicating that Ca^2+^ calmodulin enhances endophilin A1-p140Cap interaction during the initial phase of LTP.

**Figure 3.**
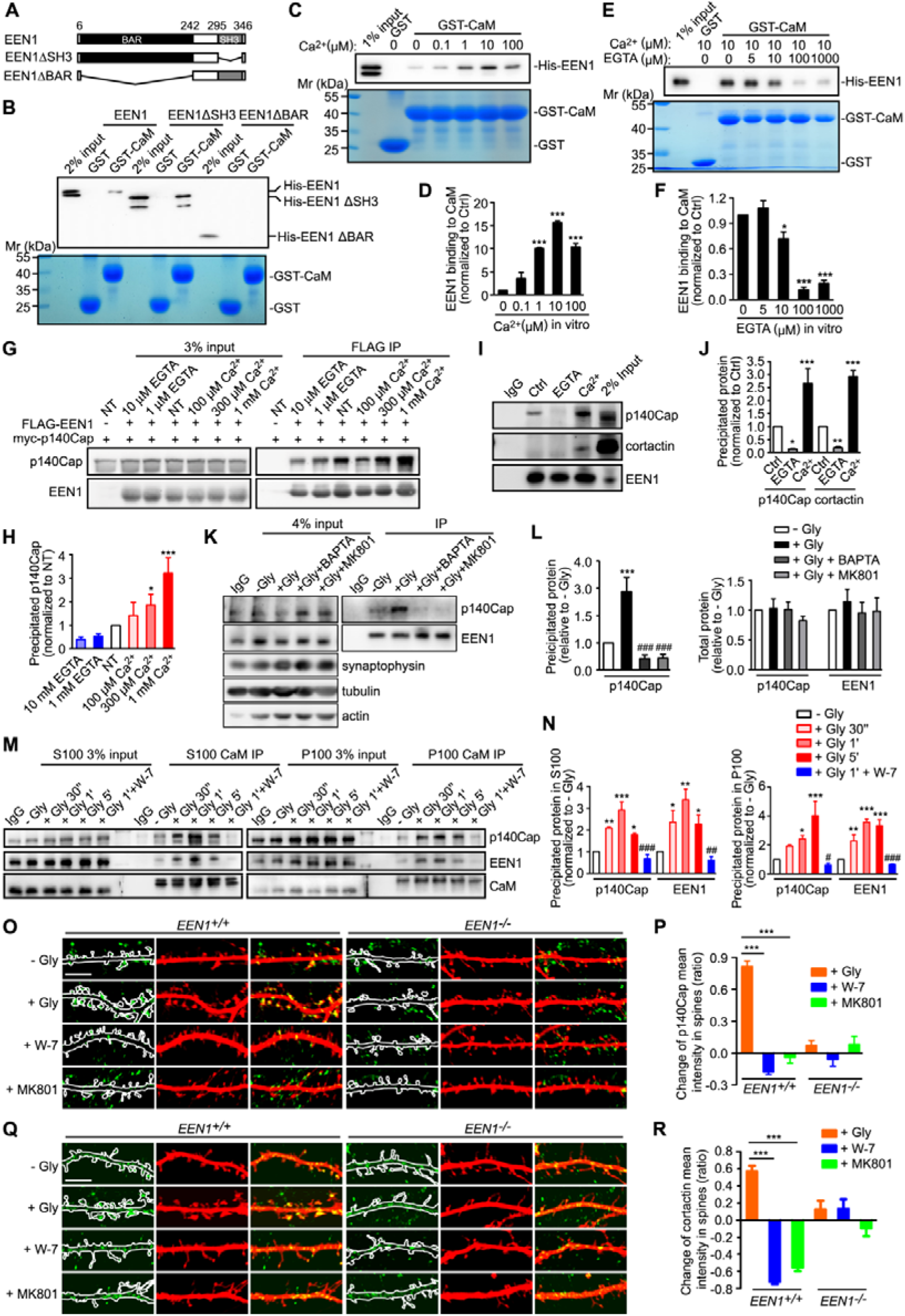
Ca^2+^/calmodulin enhances the interaction between endophilin A1 and p140Cap. (**A**) Diagram showing the domain structure and fragments of endophilin A1 used in this study. (**B**) Binding of His-tagged endophilin A1 full length, ΔSH3 and ΔBAR fragments to GST-tagged calmodulin (CaM) in GST-pull down assay. (**C**) Effect of Ca^2+^ on binding of endophilin A1 to calmodulin in GST-pull down assay. (**D**) Quantification of Ca^2+^-regulated endophilin A1 binding to calmodulin in (**C**). N = 3 independent experiments. (**E**) Effect of EGTA on binding of endophilin A1 to calmodulin in GST-pull down assay. (**F**) Quantification of endophilin A1 binding to calmodulin in E. N = 5. (**G**) Effect of EGTA or Ca^2+^ on binding of endophilin A1 to p140Cap in transiently transfected HEK293 cells in Flag IP assay. (**H**) Quantification of endophilin A1 binding to p140Cap in (**G**). N = 4. (**I**) Effect of EGTA or Ca^2+^ on binding of endophilin A1 to p140Cap in cultured neurons. Endogeneous IP assay was performed from lysates of mouse neurons with antibodies to endophilin A1. (**J**) Quantification of endophilin A1 binding to p140Cap in (**I**). N = 3. (**K**) Effect of BAPTA or MK801 on the binding of endophilin A1 to p140Cap in neurons upon cLTP induction. (**L**) Quantification of endophilin A1 binding to p140Cap and total protein levels of endophilin A1 or p140Cap in (**K**). N = 5. (**M**) Effect of W-7 on interactions between calmodulin and endophilin A1/p140Cap upon cLTP induction. DIV16 neurons were collected and the cytosolic (S100) and membrane (P100) fractions were used for immunoisolation with antibodies to calmodulin. (**N**) Quantification of endophilin A1 and p140Cap precipitated by antibodies to calmodulin in (**N**). N = 3. (**O**) Effects of W-7 and MK801 on the amount of p140Cap in spines upon LTP induction. Neurons transfected with pLL3.7-DsRed on DIV12 were pre-treated with DMSO, W-7 or MK801 and chemically induced LTP on DIV16. Neurons were fixed 1 min after glycine application, immunostained with antibodies to p140Cap (green) and imaged by confocal microscopy. Bar = 5 μm. (**P**) Quantification of changes in p140Cap mean intensity in spines as compared with the control (- Gly) group in (**O**). N ≥ 12 cells, n ≥ 464 spines per group. (**Q**) Same as (**O**) except that neurons were immunostained with antibodies to cortactin. Bar = 5 μm. (**R**) Quantification of changes in cortactin mean intensity in spines as compared with the control (- Gly) group in (**Q**). N ≥ 11 cells, n ≥ 431 spines per group. Data represent mean ± SEM in (**D**), (**F**), (**H**), (**J**), (**L**), (**N**), (**P**) and (**R**). * *p* < 0.05, ** *p* < 0.01, *** *p* < 0.001, when compared with Ctrl, NT or – Gly. ### *p* < 0.001 in (**L**), when compared with + Gly. # *p* < 0.05, ## *p* < 0.01, ### *p* < 0.001 in (**N**), when compared to + Gly 1’.

As regulator of actin remodeling, p140Cap recruits cortactin to drive Arp2/3-mediated branched actin polymerization (Jaworski et al., 2009; Schnoor et al., 2018; Yang et al., 2015). Indeed, co-IP of not only p140Cap but also cortactin by antibodies against endophilin A1 was enhanced by Ca^2+^ (Figure 3I and 3J). To test the idea that endophilin A1 functions via p140Cap and cortactin during the initial phase of sLTP, first we determined whether they are recruited to dendritic spines in an LTP- and endophilin A1-dependent manner. Quantitative analysis of immunofluorescence confocal images indicated that enrichment of p140Cap and cortactin in spines upon LTP induction requires not only endophilin A1 but also activities of calmodulin and NMDAR (Figure 3O-3R).

Next, to determine whether Ca^2+^/calmodulin promotes actin polymerization via the endophilin A1◻p140Cap pathway, we generated a calmodulin-binding deficient mutant of endophilin A1 (I154AL158A, DM) that has much lower affinity for calmodulin than the WT protein (Figure 4A and 4B; and Figure 4-figure supplement 1), and tested whether it can rescue the sLTP phenotype of *EEN1* KO neurons. Indeed, WT but not the DM mutant of endophilin A1 restored the rapid increase in spine size and F-actin content in spines upon LTP induction (Figure 4C-4F). Moreover, Y343A, the p140Cap-binding deficient mutant of endophilin A1 (Yang et al., 2015), also failed to rescue the sLTP phenotype of *EEN1* KO neurons (Figure 4G-4I). Together, these data indicate that Ca^2+^/calmodulin enhances the recruitment of p140Cap by endophilin A1 to promote actin polymerization during the initial phase of sLTP.

**Figure 4.**
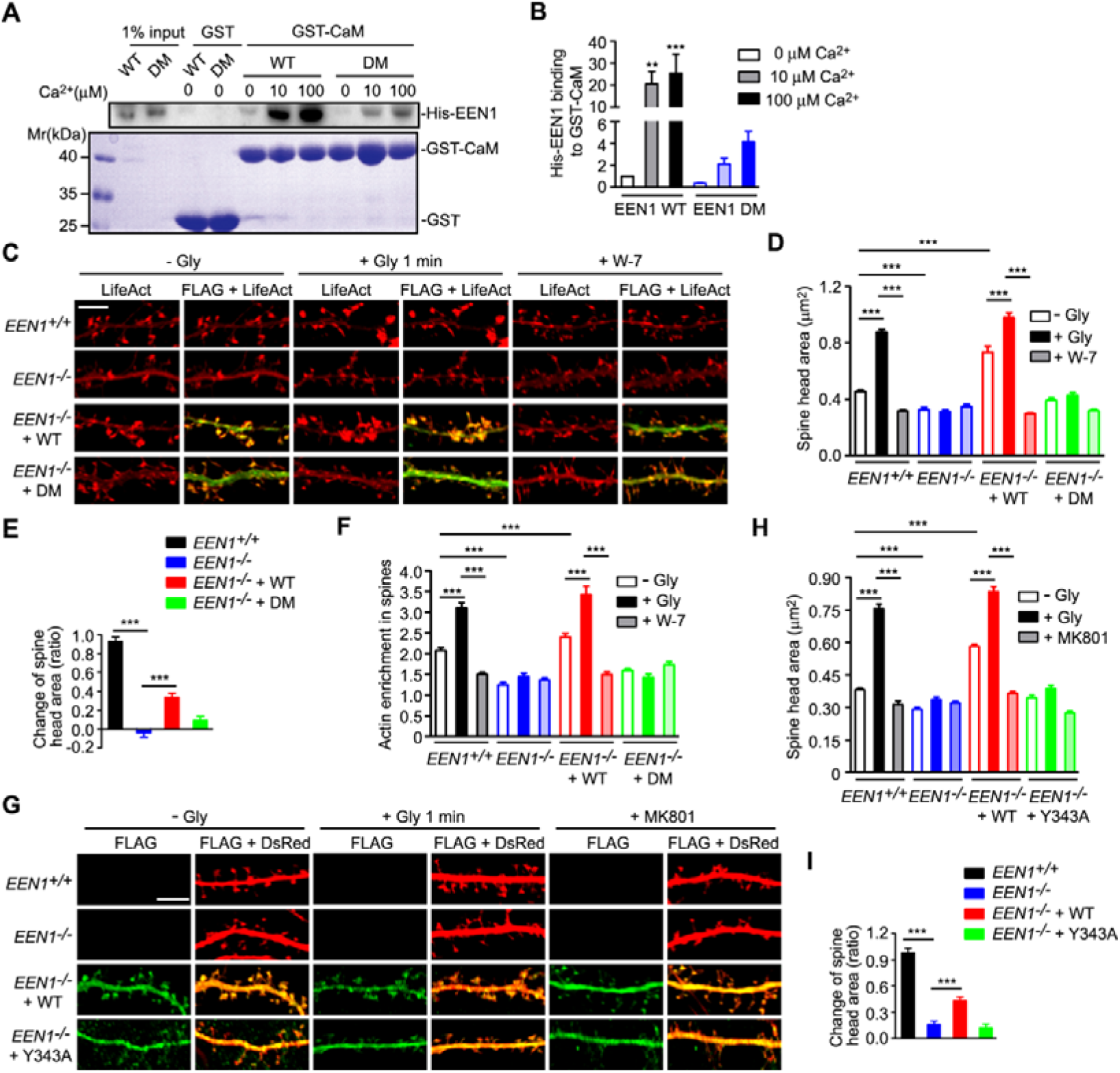
Ca^2+^/calmodulin promotes actin polymerization in spines via the endophilin A1-p140Cap pathway. (**A**) Effect of Ca^2+^ on binding of EEN1 double mutant (DM) to calmodulin in GST-pull down assay. (**B**) Quantification of EEN1 DM binding to GST-CaM as compared with WT in (**A**). Y axis shows as two segments. N = 4, *** *p* < 0.001. (**C**) Cultured *EEN1*^*+/+*^ and *EEN1*^*−/−*^ hippocampal neurons co-transfected with LifeAct-mCherry and pCMV-Tag2B, and *EEN1*^*−/−*^ hippocampal neurons co-transfected with LifeAct-mCherry and pCMV-Tag2B-EEN1 WT or pCMV-Tag2B-EEN1 DM on DIV12 were pre-treated with DMSO or W-7 and chemically induced LTP (cLTP) on DIV16. Neurons were fixed 1 min after glycine application and imaged by confocal microscopy. Bar = 5 μm. (**D, E**) Quantification of spine size and changes in spine size in (**C**). N ≥ 12 neurons, n ≥ 450 spines per group. (**F**) Quantification of F-actin enrichment in spines in (**C**). N ≥ 12 neurons, n ≥ 450 spines per group. (**G**) Cultured *EEN1*^*+/+*^ and *EEN1*^*−/−*^ hippocampal neurons co-transfected with pLL3.7-DsRed and pCMV-Tag2B, and *EEN1*^*−/−*^ hippocampal neurons co-transfected with pLL3.7-DsRed and pCMV-Tag2B-EEN1 WT or pCMV-Tag2B-EEN1 Y343A on DIV12 were pre-treated with DMSO or MK801 and chemically induced LTP (cLTP) on DIV16. Neurons were fixed 1 min after glycine application and imaged by confocal microscopy. Bar = 5 μm. (**H, I**) Quantification of spine size and changes in spine size in (**G**). N ≥ 12 neurons, n ≥ 502 spines per group. Data represent mean ± SEM in (**B**), (**D**), (**E**), (**F**), (**H**), and (**I**). ** *p* < 0.01, *** *p* < 0.001 in (**B**) when compared with 0 μm Ca^2+^.

### Ca^2+^/calmodulin-Dependent Increase in Plasma Membrane-Associated Endophilin A1 Nanodomains Correlates with Spine Size During the Initial Phase of sLTP

Single protein tracking and super-resolution imaging revealed dynamic changes in the nanoscale organization of branched F-actin regulators in spines during synaptic plasticity (Chazeau et al., 2014). Intriguingly, although the interaction between endophilin A1 and p140Cap is required for sLTP, overexpression of p140Cap could not rescue plasticity phenotypes of *EEN1* KO neurons (Yang et al., 2018), suggesting the existence of spatiotemporal regulation of their interaction during LTP induction. To this end, we analyzed the sub-spine localization of endophilin A1 by immunofluorescence staining and 3D-structured illumination microscopy (3D-SIM). Interestingly, in spines endophilin A1 was organized into nanoscale objects (mean area 0.014 μm^2^, referred to as nanodomains) which did not overlap with PSD95, marker for the postsynaptic density (PSD) structure (Figure 5-figure supplement 1A). Quantitative analysis revealed an NMDAR-dependent increase in the number of endophilin A1 nanodomains in spines undergoing sLTP (Figure 5-figure supplement 1A-1C). Moreover, the size of spine head correlated with the number but not the area of endophilin A1 nanodomains (F Figure 5-figure supplement 1E and 1F). In contrast, the number of endophilin A1 nanodomains did not correlate with the size of the PSD95-labeled PSD structures (Figure 5-figure supplement 1D and 1G). Further, treatment of hippocampal neurons with inhibitor of calmodulin but not CaMKII abolished the increase in not only spine size but also the number of endophilin A1 puncta in spines (Figure 5-figure supplement 1H-1J). These data together suggest that the calmodulin-regulated subsynaptic localization of endophilin A1 is required for spine enlargement.

As endophilin A1 contains the positive membrane curvature-sensing and binding N-BAR domain (Gallop et al., 2006) that enables it associate with invaginated plasma membrane, next we investigated whether its association with the spine plasma membrane is also regulated in the initial phase of sLTP. To distinguish plasma membrane-localized endophilin A1 from those localized to intracellular structures, we permeabilized cell membrane with the mild detergent saponin to limit access of antibodies to the cytosolic leaflet of the plasma membrane. Indeed, most endophilin A1 nanodomains were plasma membrane-localized (Figure 5A). Moreover, there was an increase in the number of plasma membrane-localized endophilin A1 nanodomains as early as 1 minute after LTP induction (Figure 5B). Further, the number but not the area of the nanodomains correlated with the size of spine head (Figure 5B-5D). As endophilin A1 functions via p140Cap and its downstream effector cortactin (Yang et al., 2015) to activate Arp2/3 which in turn induces branched actin polymerization (Uruno et al., 2001; Weaver et al., 2001), these data suggest that upon LTP induction, endophilin A1 localizes to the periphery of spine head and promotes local actin polymerization underneath the plasma membrane.

**Figure 5.**
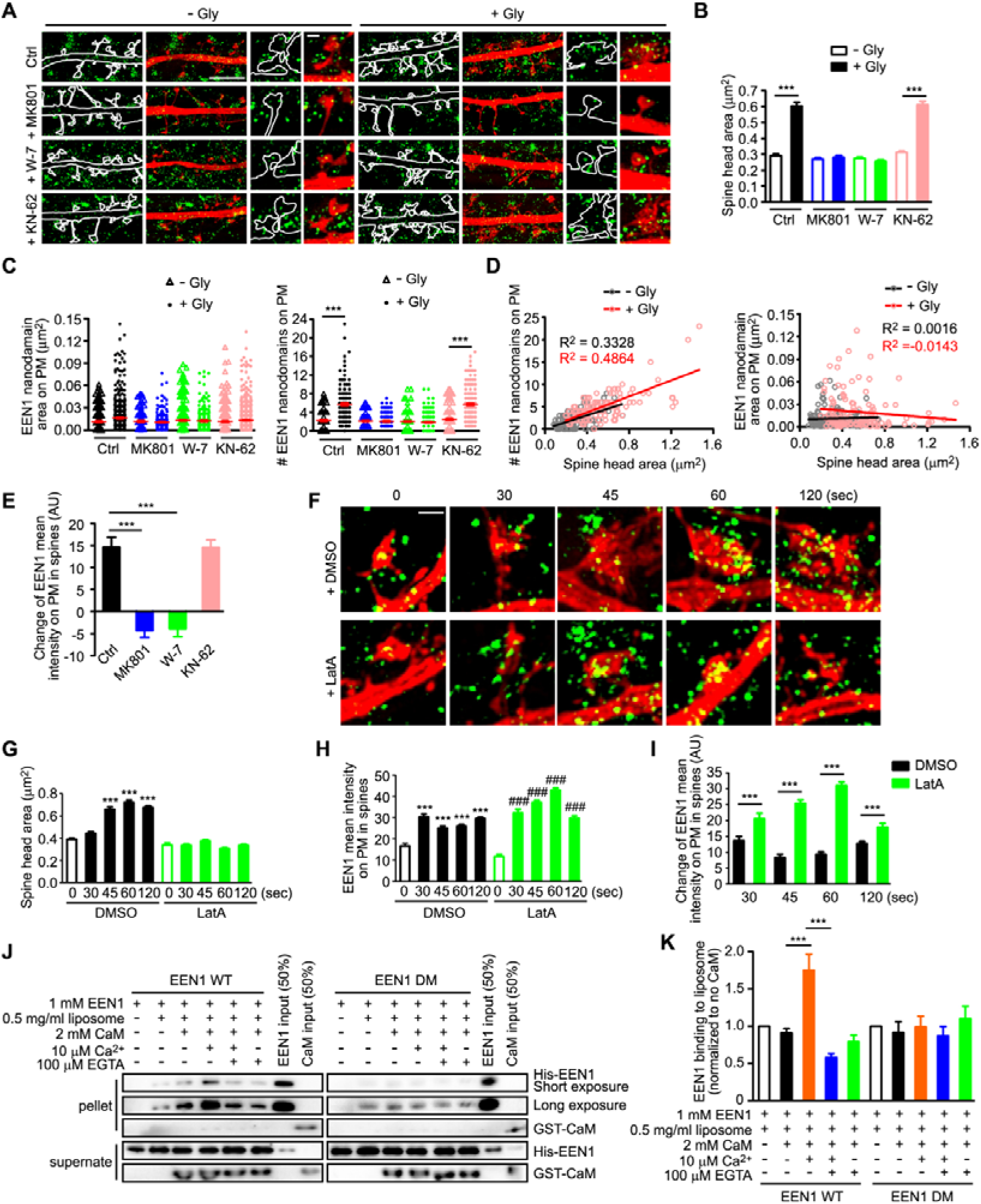
Ca^2+^/calmodulin-enhanced plasma membrane association of endophilin A1 correlates with spine expansion. (**A**) DIV16 neurons were pre-treated with DMSO, W-7 or KN-62 and chemically induced LTP. Neurons were fixed 1 min after glycine application. Plasma membrane (PM)-localized endophilin A1 was immunostained and imaged by 3D-SIM. Spines were outlined manually. Bar = 4 μm in left and center panels and 500 nm in magnified images in right panels. (**B**) Quantification of spine size in (**A**). (**C**) Quantification of the area and number of PM-localized endophilin A1 nanodomains in spines in (**A**). (**D**) Scatterplot of the number or area of PM-localized endophilin A1 nanodomains versus the size of spine head for control (- Gly, n = 278 spines) and cLTP (+ Gly, n = 254 spines) groups with linear fits. (**E**) Quantification of changes in the mean intensity of PM-localized endophilin A1 in spines. (**F**) DIV16 neurons were pre-treated with DMSO or LatA and chemically induced LTP. Neurons were fixed at 30 s, 45 s, 60 s or 120 s after glycine application. PM-localized endophilin A1 was stained and imaged by 3D-SIM. Bar = 500 nm. (**G**) Quantification of spine size in (**F**). (**H**) Quantification of the mean intensity of PM-localized endophilin A1 in spines in (**F**). (**I**) Quantification of changes in the mean intensity of PM-localized endophilin A1 in spines in (**F**). (**J**) Effect of Ca^2+^/calmodulin on binding of EEN1 WT or DM protein to membrane in liposome sedimentation assay. (**K**) Quantification of endophilin A1 binding to liposome in (**J**). N = 7. Data represent mean ± SEM in B-E, G-I, and K. *** *p* < 0.001, when compared with – Gly in (**G**) and (**H**). ### *p* < 0.001, when compared with – Gly + LatA treatment in (**H**). N ≥ 10 neurons, n ≥ 254 spines per group in (**B-E**), and N ≥ 10 neurons, n ≥ 241 spines per group in (**G-I**).

Given that sub-spine accumulation of endophilin A1 requires activation of NMDAR and calmodulin (Figure 5-figure supplement 1), we reasoned that upon NMDAR-mediated Ca^2+^ influx, binding of Ca^2+^-activated calmodulin to endophilin A1 not only enhances its interaction with p140Cap, but also facilitates its association with the invaginated plasma membrane. Indeed, inhibition of NMDAR or calmodulin, not CaMKII, abolished glycine-induced increase in the number of plasma membrane-localized endophilin A1 nanodomains (Figure 5A, 5C and 5E). Notably, although latrunculin A treatment inhibited spine enlargement, it had no effect on LTP-induced increase in plasma membrane-localized endophilin A1 (Figure 5F-5I), indicating that Ca^2+^/calmodulin-enhanced association of endophilin A1 with the plasma membrane precedes actin polymerization during sLTP initiation.

To corroborate that Ca^2+^/calmodulin regulates the association of endophilin A1 with the plasma membrane, we performed *in vitro* liposome sedimentation assays and found that, whilst calmodulin alone did not change the membrane-binding capacity of WT endophilin A1, Ca^2+^/calmodulin enhanced it significantly (Figure 5J and 5K). In contrast, Ca^2+^/calmodulin had no effect on the membrane association ability of the calmodulin-binding deficient DM mutant of endophilin A1 (Figure 5J and 5K). Collectively, these data indicate that Ca^2+^/calmodulin enhances association of endophilin A1 with the plasma membrane of dendritic spines in the initial phase of sLTP.

### Endophilin A1 Promotes Branched Actin Polymerization Underneath the Plasma Membrane

Based on the findings that endophilin A1 accumulates into spine plasma membrane-associated nanodomains and recruits p140Cap and cortactin in the initial phase of sLTP, we further reasoned that it might transduce the Ca^2+^ signals instantaneously to enable plasma membrane expansion of spines by promoting branched actin polymerization. If it is true, we should be able to detect more endophilin A1 associated with both membranes and the actin cytoskeleton upon LTP induction. To test this possibility, first we analyzed subcellular fractions of mouse hippocampi from animals subjected to fear conditioning, a physiological learning paradigm associated with synaptic plasticity. Indeed, compared with naïve mice, we detected significant increase in the amount of endophilin A1 and p140Cap in both membrane and cytoskeletal fractions from trained animals (Figure 6A and 6B). Consistently, although the levels of either protein remained unchanged (Figure 6C and 6D), their association with membrane and cytoskeleton fractions also increased in dissociated cultured hippocampal neurons in the initial phase of cLTP (Figure 6E and 6F). Moreover, the enhanced association of endophilin A1 and p140Cap with both membrane and cytoskeleton was inhibited by W-7 but not KN-62 (Figure 6E and 6F), indicating that calmodulin is the immediate upstream regulator of their subcellular redistribution. Notably, subcellular distribution of endophilin A2, another member of the endophilin A family, was not affected by either neural activity or Ca^2+^/calmodulin (Figure 6A-6F).

**Figure 6.**
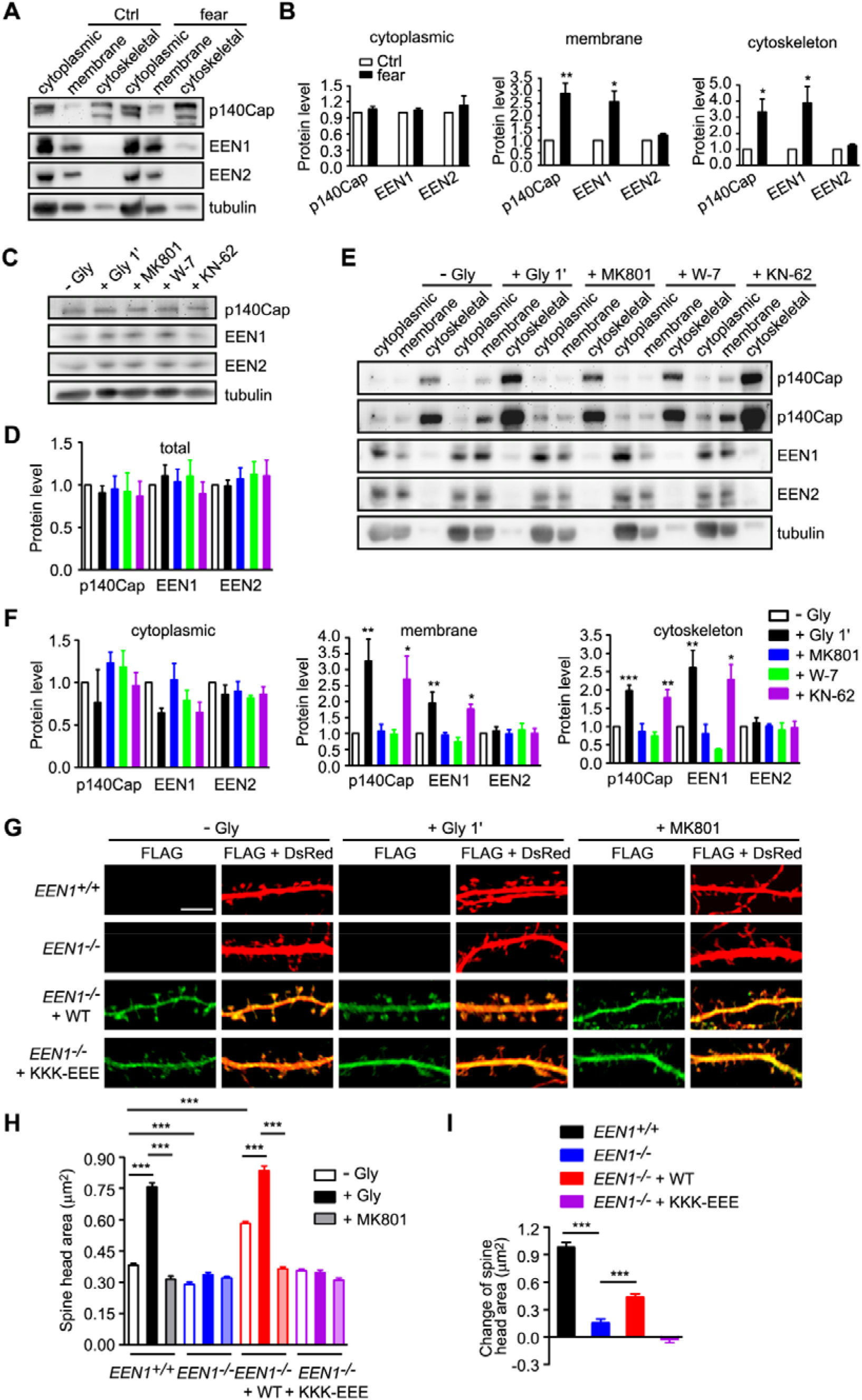
Ca^2+^/calmodulin regulates the association of endophilin A1 and p140Cap with both membrane and cytoskeleton upon LTP induction. (**A**) Immunoblotting of subcellular fractionations of mouse hippocampi from animals subjected to fear training. (**B**) Quantification of protein levels of p140Cap, endophilin A1 or endophilin A2 in cytoplasmic, membrane and cytoskeleton fractions in (**A**). N = 6 animals. (**C**) Cultured neurons were pre-treated with DMSO, 10 μM MK801, W-7 or KN-62 and chemically induced LTP on DIV16. Cell lysates were prepared 1 min after glycine application and subjected to SDS-PAGE and immunoblotting. (**D**) Quantification of total protein levels in (**C**). N = 5. (**E**) Effects of MK801, W-7 and KN-62 on subcellular distribution of protein levels. Neurons were pre-treated with DMSO, MK801, W-7 or KN-62 and chemically induced LTP on DIV16. Cells were collected and subjected to subcellular fractionation 1 min after glycine application. (**F**) Quantification of protein levels in subcellular fractions in (**E**). N = 5. (**G**) *EEN1*^*+/+*^ and *EEN1*^*−/−*^ hippocampal neurons co-transfected with pLL3.7-DsRed and pCMV-Tag2B, and *EEN1*^*−/−*^ hippocampal neurons co-transfected with pLL3.7-DsRed and pCMV-Tag2B-EEN1 WT or pCMV-Tag2B-EEN1 KKK-EEE on DIV12 were pre-treated with DMSO or MK801 and chemically induced LTP on DIV16. Neurons were fixed 1 min after glycine application and imaged by confocal microscopy. Bar = 5 μm. (**H, I**) Quantification of spine size or changes in spine size in (**G**). N ≥ 12 neurons, n ≥ 482 spines per group. Data represented are mean ± SEM in (**B**), (**D**), (**F**), (**H**) and (**I**). * *p* < 0.05, ** *p* < 0.01, *** *p* < 0.001 when compared to Ctrl or – Gly in (**B**) and (**F**).

We then determined whether the membrane-binding capacity of endophilin A1 is required for the rapid spine enlargement of neurons undergoing sLTP. Compared with WT, the membrane-binding deficient mutant of endophilin A1 (KKK-EEE) (Gallop et al., 2006) was unable to rescue the deficit in acute spine enlargement of glycine-treated *EEN1* KO neurons (Figure 6G-6I). All together, these data indicate that rapid enlargement of spines in the initial phase of sLTP requires Ca^2+^/calmodulin-dependent enhancement of not only endophilin A1-p140Cap interaction but also the association of endophilin A1 with membrane.

Since the small size of dendritic spines and the wide distribution of F-actin and actin polymerization regulators in spines prevent us to better visualize the spatiotemporal relationship between plasma membrane association of endophilin A1 and its effectors, we tested whether there are changes in the subcellular distribution of endophilin A1and Arp2/3 in response to Ca^2+^/calmodulin using HeLa cell as a heterologous model system. In HeLa cells ectopically co-expressing endophilin A1 and p140Cap, upon Ca^2+^ influx induced by the calcium ionophore ionomycin, we detected enrichment of endophilin A1 signals underneath the plasma membrane by confocal microscopy (Figure 6-figure supplement 1A). Moreover, preincubation with W-7 but not latrunculin or CK-666 abolished ionomycin-induced endophilin A1 recruitment to the cell periphery (Figure 6-figure supplement 1A), indicating that Ca^2+^-activated calmodulin directly regulates accumulation of endophilin A1 at the plasma membrane. Further, although the strong intrinsic signals for cortical actin did not allow us to quantify changes in F-actin content underneath the plasma membrane, the Arp2/3 complex (labeled with fluorescently tagged Arp1b) was also enriched in the cell periphery upon ionomycin application (Figure 6-figure supplement 1B). In contrast, ionomycin treatment failed to cause recruitment of either the membrane-binding (KKK-EEE) or the calmodulin-binding deficient mutant of endophilin A1 to the cell periphery (Figure 6-figure supplement 1B). Intriguingly, although ionomycin treatment increased plasma membrane association of the p140Cap-binding mutant Y343A, no enrichment of Arp1b signals at the cell periphery was observed (Figure 6-figure supplement 1B), indicating that recruitment of the Arp2/3 complex underneath the plasma membrane requires the interaction between endophilin A1 and p140Cap. Collectively, these results support that Ca^2+^/calmodulin enhances association of endophilin A1 with the plasma membrane and promotes branched actin polymerization in spines.

### Ca^2+^/calmodulin-Regulated Functions of Endophilin A1 is Required for LTP and Long-term Memory

It was proposed that spine enlargement enables formation of a stable F-actin:cofilin complex that serves as a synaptic tag to capture postsynaptic constituent proteins for maintenance and consolidation of the potentiated state (Bosch et al., 2014). To determine the functional significance of endophilin A1-mediated structural plasticity, next we tested whether the molecular functions of endophilin A1 in structural plasticity are also required for LTP by electrophysiological analyses. To this end, we performed molecular replacement in a small subset of neurons by injection of the hippocampal CA1 region of *EEN1*^fl/fl^ mice with adeno-associated viral vectors encoding the Cre recombinase (AAV-mCherry-Cre) together with those encoding either WT or mutant endophilin A1 (AAV-EGFP-2A-EEN1 WT, Y343A, KKK-EEE or DM) at postnatal day 0 (P0), and induced LTP in Schaffer-collateral synapses by double patch whole-cell recording of non-infected (control, *EEN1*^fl/fl^) and virus-infected CA1 pyramidal cells in acute slices from virus-injected animals at P14-21. In line with our previous findings (Yang et al., 2018), Cre-mediated knockout of *EEN1* caused significant impairment of LTP, which was fully rescued by re-expression of WT endophilin A1 (Figure 7A, 7B and 7F). In contrast, none of the mutants could rescue the magnitude or the maintenance of LTP in *EEN1*^−/−^ neurons (Figure 7C-7F), indicating that indeed, molecular functions of endophilin A1 to initiate sLTP is required for expression and stabilization of synaptic potentiation.

**Figure 7.**
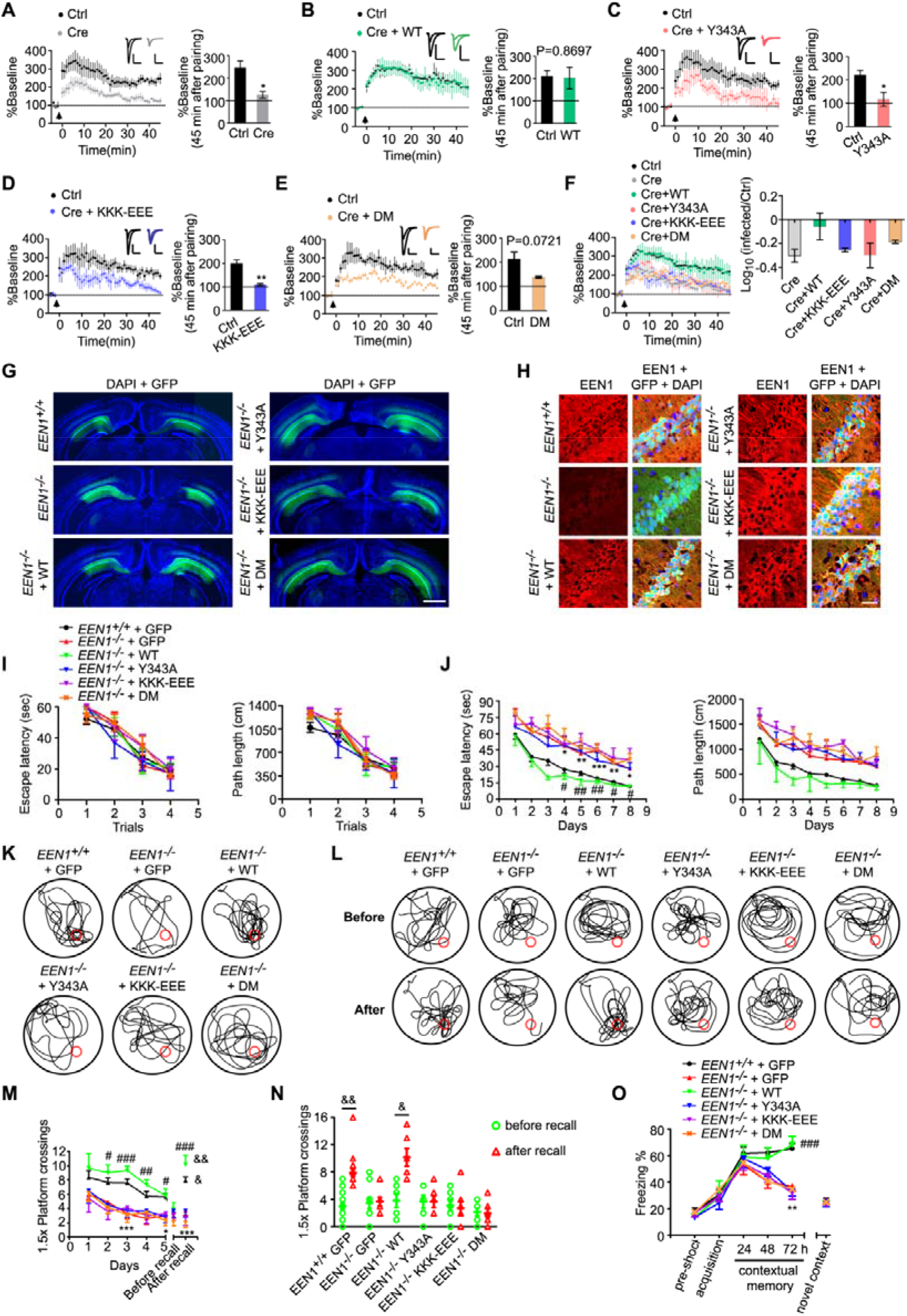
The calmodulin-, membrane- and p140Cap-binding capacities of endophilin A1 are required for LTP and long-term memory. (**A-F**) Rescue of the LTP impairment phenotype in *EEN1* KO neurons by EEN1 WT and mutants. AAV viruses expressing the Cre recombinase (AAV-mCherry-2A-Cre) and GFP (AAV-EGFP) or Cre, GFP and EEN1 WT or mutant (AAV-EGFP-2A-EEN1) were stereotaxically injected into the CA1 regions of EEN1^fl/fl^ mice at P0. Acute slices of hippocampi were prepared on P14-P21 for dual recording analysis of LTP. Shown are pairwise comparisons of LTP in non-infected (Ctrl) and infected neurons of the same slice. For Cre-expressing neurons (Cre) versus Ctrl in (**A**), 6 recording pairs from three mice, marked as N = 6/3, were analyzed. N = 10/5 in (**B**), N = 5/4 in (**C**), N = 5/3 in (**D**) and N = 4/3 in (**E**). Bar graphs shown percentage of baseline at 45 min after pairing. A summary of rescue effects of EEN1 WT and mutants is shown in (**F**). Data represent mean◻±◻SEM. * *p* < 0.05, ** *p* <0.01. (**G**) AAV virus was stereotaxically injected into the CA1 regions of *EEN1*^*+/+*^ to express GFP alone, or *EEN1*^*−/−*^ mice to express GFP alone, EEN1 WT, KKK-EEE, Y343A or DM and GFP. Shown are images of GFP fluorescent signals and DAPI labeling of nuclei captured by fluorescence microscopy. Bar = 1 mm. (**H**) Immunofluorescence staining of endophilin A1 in CA1 neurons of brain slices from mice in (**G**). Bar = 50 μm. (**I-N**) Morris water maze test of AAV-injected mice. Shown are escape latency or distance travelled before escaping to the platform in the visible platform training (**I**), escape latency and distance travelled before escaping to the platform in the invisible platform training (**J**), the swim trace in probe trial 3 and recall following training once again one month after training (**K, L**), number of crossings within the 1.5× platform area in 5-day probe trial and recall test (**M**), and number of crossings within the 1.5× platform area before and after recall (**N**). (**O**) Freezing behavior in AAV-injected *EEN1*^*+/+*^ or *EEN1*^*−/−*^ mice subjected to contextual fear conditioning. Data represent mean◻±◻SEM in H-O (15 *EEN1*^*+/+*^ + GFP, 10 *EEN1*^*−/−*^ + GFP, 6 *EEN1*^*−/−*^ + EEN1, 6 *EEN1*^*−/−*^ + Y343A, 7 *EEN1*^*−/−*^ + KKK-EEE, 7 *EEN1*^*−/−*^ + DM), * *p* < 0.05, ** *p* <0.01, *** *p* < 0.001 when compared with *EEN1*^*+/+*^ + GFP, # *p* < 0.05, ## *p* < 0.01, ### *p* < 0.001 when compared with *EEN1*^*−/−*^ + GFP; & *p* < 0.05, && *p* < 0.01 when compared with before recall.

Finally, we determined the physiological significance of the mechanistic roles of endophilin A1 in LTP by testing whether endophilin A1 mutants can rescue the learning and memory deficits in *EEN1* KO mice. Indeed, whilst AAV-mediated expression of WT endophilin A1 in the CA1 region restored the long-term memory in KO mice in both Morris water maze and fear conditioning tests, neither of the three mutants ameliorated the phenotypes (Figure 7G-O). Collectively these data indicate that in CA1 pyramidal cells, not only the p140Cap-binding, but also the calmodulin-interaction and membrane association capacities of endophilin A1 are required for long-term synaptic potentiation and memory.

## Discussion

The temporal phases of sLTP include initiation of spine expansion (≤ 1 min after LTP stimulation, initial phase), transient (early phase) and sustained spine enlargement (late phase) (Harvey and Svoboda, 2007; Matsuzaki et al., 2004). Although actin polymerization is essential for sLTP (Matsuzaki et al., 2004; Obashi et al., 2019), little is known about the relationship between actin remodeling and membrane dynamics during the initial phase due to limited spatiotemporal resolution of light microscopy. Moreover, although imaging studies have revealed subspine organization and dynamics of F-actin pools as well as nanoscale segregation of branched F-actin regulators in dendritic spines and suggested that their localization might be spatially and temporally controlled during activity-dependent morphological changes (Chazeau et al., 2014; Honkura et al., 2008), the precise sequence of molecular events leading to rapid structural expansion of spines remains to be defined. In this work, we uncover a novel mechanism for initiation of sLTP. We show that in direct response to Ca^2+^/calmodulin, endophilin A1 drives acute spine enlargement in NMDAR-mediated sLTP by localizing to spine plasma membrane and recruiting p140Cap to promote branched actin polymerization (Figure 8).

**Figure 8.**
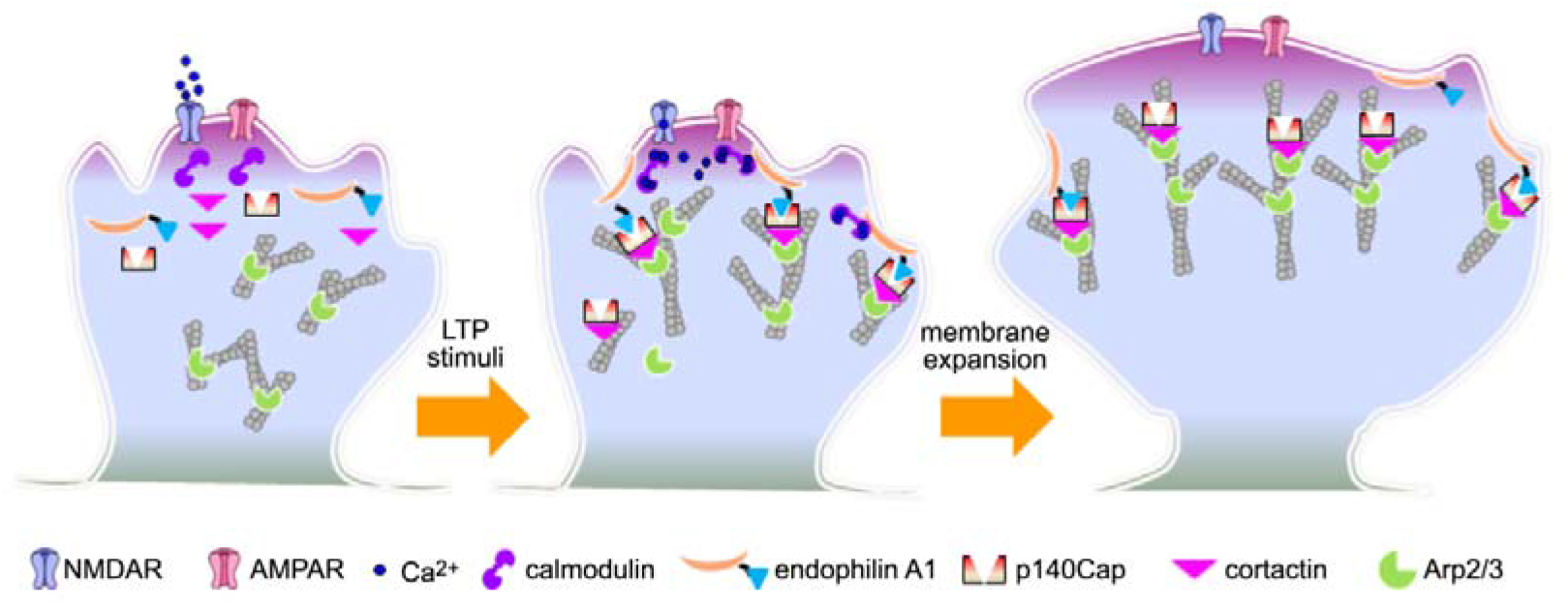
Model for endophilin A1-mediated initial expansion of spine head in sLTP. LTP stimuli induce NMDAR-mediated Ca^2+^ influx into dendritic spines and activation of calmodulin. Ca^2+^/calmodulin interacts directly with endophilin A1 and enhance its binding to both the plasma membrane and p140Cap. As a result, the plasma membrane-associated endophilin A1 recruits p140Cap, which in turn recruits cortactin to promote branched actin polymerization underneath the plasma membrane, generating propulsive force for rapid structural expansion of spine head during the initial phase of sLTP.

Endophilin A2 and A3, two other members of the endophilin A family, have been found to interact with the immediate early protein Arc/Arg3.1 to accelerate AMPAR endocytosis and might contribute to late-phase synaptic plasticity (Chowdhury et al., 2006). The impairment of structural and functional plasticity of potentiated dendritic spines caused by ablation of endophilin A1 cannot be rescued by overexpression of endophilin A2 or A3 (Yang et al., 2018). Although both endophilin A1 and A2 interact with calmodulin (Myers et al., 2016) (and Figure 3, this study), only endophilin A1, but not A2 or A3, binds to and recruits p140Cap to dendritic spines to promote actin polymerization (Yang et al., 2015). Moreover, *in vivo* and *in vitro* LTP stimuli induce increased association of endophilin A1, but not endophilin A2, with both membrane and cytoskeleton (Figure 6, this study). Together these findings indicate that different interaction partners for the endophilin A family members confer them distinct mechanistic roles in the induction and expression of synaptic plasticity in dendritic spines.

Consistent with previous reports (Yang et al., 2008), we found that SNARE-mediated membrane fusion is not required for rapid spine expansion in the initial phase of sLTP (Figure 2). Upon LTP induction, Ca^2+^ influx through the NMDAR ion channel activates calmodulin, which in turn activates CaMKII and its downstream signaling cascades to trigger various subcellular events including actin remodeling (Chazeau et al., 2014). Previous live imaging studies have revealed calmodulin-dependent formation of an F-actin pool that associates with spine enlargement (hence referred to as “enlargement pool”) (Honkura et al., 2008). Although the observation that the membrane ruffling of the spine head synchronizes with the enlargement pool of F-actin has prompted the authors to conclude that spine enlargement is induced by the propulsive force generated by calmodulin-regulated actin polymerization (Honkura et al., 2008), the mechanistic link between Ca^2+^/calmodulin and actin polymerization was still missing. In this study, we demonstrate that membrane-associated endophilin A1 is the direct molecular target of Ca^2+^/calmodulin, which enhances both its membrane association and its interaction with the downstream effector p140Cap (Figure 3 and 5). Our data further indicate that coordination of the membrane-association and p140Cap-binding capacities of endophilin A1 provides the protrusive force for rapid structural remodeling of dendritic spines by promoting actin polymerization underneath the plasma membrane (Figure 4, 6, 7 and 8), which is also in good agreement with most recent studies that the interplay between membrane tension and branched actin polymerization could produce membrane deformations (Simon et al., 2019).

Most recent studies revealed that although Rac1 activity is not required for the initial spine expansion induced by glutamate uncaging, formation of a “reciprocally activating kinase-effector complex” (RAKEC) between CaMKII, and Tiam1, a guanine exchange factor for Rac (RacGEF), converts the transient Ca^2+^ signal triggered by LTP induction into a persistent kinase signal required for the maintenance of sLTP (Saneyoshi et al., 2019). Given the spinous dynamic reorganization of nanoscale distribution of various F-actin regulators that are downstream of Ca^2+^/calmodulin and CaMKII (Chazeau and Giannone, 2016; Chazeau et al., 2014), including the F-actin severing protein cofilin (Noguchi et al., 2016), there might be crosstalk between Ca^2+^/calmodulin-dependent, endophilin A1-mediated actin polymerization and other Ca^2+^/calmodulin and/or CaMKII effector-regulated pathways (e.g., Rac1 and ADF/cofilin), which enables spatiotemporally controlled actin reorganization during the initial phase and/or the transition to the early phase of sLTP.

Copine-6, a Ca^2+^ sensor, relocalizes from the cytosol of dendrites to lipid raft-enriched postsynaptic plasma membrane in response to NMDAR-mediated Ca^2+^ influx, and activates Rac1 to spines during synaptic potentiation (Reinhard et al., 2016). It is also required for spine structural plasticity and LTP, probably contributing to stabilization of the actin cytoskeleton by inhibiting ADF/cofilin via the Rac1-PAK-LIMK1 pathway (Reinhard et al., 2016). In addition, by keeping cortactin active to prevent branched actin filaments from severing by ADF/cofilin, the fast Ca^2+^ sensor caldendrin stabilizes an F-actin pool at the spine base required for the structural remodeling of spines in the transition from early to late phase LTP (Mikhaylova et al., 2018). In our study, while repetitive expansion and shrinkage of spine heads were also observed in *EEN1* KO neurons, the F-actin content and spine size remained constant during the initial phase of sLTP (Figure 1). Rather than stabilizing F-actin, endophilin A1 responds to Ca^2+^/calmodulin and promotes branched actin polymerization beneath the plasma membrane to achieve rapid spine enlargement. While they all function via CaMKII-independent mechanisms to regulate actin dynamics, how Colpine-6, caldendrin and endophilin A1 coordinate with each other to ensure spine potentiation and stabilization awaits further investigation.

## Acknowledgements

We thank Dr. Anbing Shi (Huazhong University of Science and Technology) for critical comments on the manuscript. This work was supported by funding from the National Natural Science Foundation of China (31530039 and 91954126 to J-J. Liu, 31571056 to Y. Yang, 91849112 to Y. S. Shi and 81901161 to J. Chen, 91754202 to Dong Li and 31670846 to C. Ma), Ministry of Science and Technology of China (2016YFA0500100 to J-J. Liu and 2019YFA0801603 to Y.S. Shi), and the Strategic Priority Research Program of Chinese Academy of Science (XDB32020100).

## Author Contributions

Conceptualization, Y.Y. and J-J.L.; Investigation, Y.Y., J.C., X.C., Di.L., S.Z., X.Y., S.D., D.W., Z.G., and S.Z.; Formal analysis, Y.Y., J.C., X.C., J.H., S.Z., and S.D.; Visualization, Y.Y., J.C., X.C., J.H., S.Z., S.D., X.L. and S.D.; Writing – Original Draft, Y.Y., J.C., S.Z., X.Y., X.L., Y.S.S., and J-J.L.; Funding Acquisition, Y.Y., J.C., Dong L., C.M., Y.S.S. and J-J.L.; Supervision, Dong.L., C.M., X.L., Y.S.S., and J-J.L.; Project Administration, J-J.L.

## Materials and Methods

### Ethics statement

All animal experiments were approved by and performed in accordance with the guidelines of the Animal Care and Use Committee of Institute of Genetics and Developmental Biology, Chinese Academy of Sciences (Approval code: AP2013003 and AP2015002), and the Animal Care and Use Committee of the Model Animal Research Center, the Host for the National Resource Center for Mutant Mice in China, Nanjing University (Approval code: AP#SY06). All animals were housed in standard mouse cages at 22-24 °C on a 12 h light/dark cycle with access to food and water freely.

### Animals

Generation of *EEN1*^fl/fl^ and *EEN1*^−/−^ mice on the C57BL/6J background was as previously described (Yang et al., 2018). Briefly, the targeting vector for *EEN1* was obtained from European Mouse Mutant Cell Repository (EuMMCR, PRPGS00060_A_A02). The endophilin A1 KO first and *EEN1*^fl/fl^ C57BL/6J mice were generated at Nanjing Model Animal Research Center of Nanjing University. Genotyping of mouse lines was performed by genomic PCR of tail prep DNA from offspring with the following primer pairs: loxPF/loxPR: 5’-CAAGGACTCCCAGAGACCTAGCATC-3’ and 5’-GAGATGGCGCAACGCAATTAAT-3’ (A PCR product of 375 base pairs in EEN1 KO first mice but none in wild-type mice).

zptF/zptR: 5’-GTAAGCGGCTCTAGCGCATGTTCT-3’ and 5’-GCAGGGGCATGTAGGTGGCTCAAC-3’ (A PCR product of 466 base pairs in WT mice, none in EEN1 KO first mice, and of 627 base pairs in *EEN1*^fl/fl^ mice).

### Constructs

The putative amino acid residues of EEN1 involved in calmodulin binding were predicated using the Binding Site Search and Analysis tool provided at the Calmodulin Target Database: http://calcium.uhnres.utoronto.ca/ctdb/ctdb/home.html. pCMV-Tag2B-EEN1 DM and pET28a(+)-EEN1 single mutants and DM (mutation I154 and L158 to A) were created by site-directed mutagenesis using pCMV-Tag2B-EEN1 and pET28a(+)-EEN1 as template, respectively. pET28a(+)-EEN1 ΔBAR (Δ aa 6-242) and pET28a(+)-EEN1 ΔSH3 (Δ aa 295-346) were subcloned from pGEX4T-1-EEN1 ΔBAR and pGEX4T-1-EEN1 ΔSH3. The pAAV-CaMKIIα-EGFP-2A-MCS-3FLAG-EEN1 or mutant of EEN1 AAV viral constructs (KKK-EEE, Y343A and DM) were generated by cloning EEN1 cDNA amplified from pCMV-Tag2B-EEN1 or mutant constructs into the pAAV-CaMKIIα-EGFP-2A-MCS-3FLAG. The bacterial expression construct for His-tagged p140Cap fragment (aa 351-1051) was generated by PCR amplification of the cDNA encoding p140Cap (aa 351-1051) and insertion into pET-28a(+). Bacterial expression constructs for calmodulin were generated by PCR amplification of cDNA for mouse calmodulin from mouse brain by RT-PCR and insertion into pGEX4T-1 and pET28a(+). Arp1b-mCherry construct was a generous gift from Drs. Na Mi and Li Yu (Tsinghua University, China). All other constructs used in this study (pCMV-Tag2B-EEN1 WT, pCMV-Tag2B-EEN1 Y343A, pCMV-Tag2B-EEN1 KKK-EEE, mGFP and LifeAct-mCherry) were described previously (Guo et al., 2018; Yang et al., 2018). Viral particles of adeno-associated virus (AAV) carrying pAOV-CaMKIIα-EGFP-2A-Cre, pAAV-CaMKIIα-EGFP-2A-MCS-3FLAG-EEN1 or mutants of EEN1 and the control construct pAAV-CaMKIIa-EGFP-2A-MCS-3FLAG were purchased from OBiO Technology (Shanghai) Corp. Ltd., (Shanghai, China).

### Antibodies

The following antibodies were obtained from commercial sources: goat anti-endophilin A1 (S-20), endophilin A2 (E-15), mouse anti-SYP (D-4), and mouse anti-cortactin (E-4) (Santa Cruz Biotechnology, Santa Cruz, CA, USA) for WB and anti-cortactin (EMD Millipore, Temecula, CA, USA) for staining; rabbit anti-endophilin A1 (Synaptic Systems GmbH, Germany); rabbit anti-Myc, mouse anti-GST, rabbit and mouse anti-GFP, rabbit and mouse anti-RFP which recognizes DsRed and mCherry (Medical & Biological Laboratories, Naka-ku Nagoya, Japan); mouse anti-M5 DYKDDDDK-Tag (Mei5 Biotechnology, Beijing, China), mouse anti-α-tubulin and mouse anti-β-actin (Sigma-Aldrich, St. Louis, MO, USA); mouse anti-His (CoWin Biosciences, Jiangsu, China), rabbit anti-calmodulin (Boster Biological Technology, Pleasanton, CA, USA) for WB and mouse anti-calmodulin (Invitrogen, Carlsbad, CA, USA) for IP; Rabbit anti-p140Cap was described previously (Yang et al., 2015). Secondary antibodies for immunofluorescence staining were from Molecular Probes (Invitrogen, Carlsbad, CA, USA).

### Cell culture, transfection and drug treatment

Primary hippocampal neurons were cultured as previously described (Yang et al., 2018). Briefly, mouse hippocampi were dissected from P0 C57BL/6J mice, dissociated with 0.125% trypsin in Hank’s balanced salt solution without Ca^2+^ and Mg^2+^ at 37°C for 15 min, triturated in DMEM, 10% F12, and 10% FBS (Gibco, Carlsbad, CA, USA). Hippocampal neurons were plated on poly-D-lysine-coated coverslips in 24-well plates or 30-mm dishes at a density of 2.5-3.0 × 10^4^ cells/well in 24-well plate or 1.0-1.2 × 10^5^ cells/35 mm dish. The medium was replaced with the serum-free Neurobasal A (NB-A) media supplemented with 2% B27 supplement, GlutaMAX (Gibco, Carlsbad, CA, USA) and 0.3% glucose 4 h after plating. Half of the media were replaced every 3 days until use.

For neuronal morphology and immunofluorescence staining, neuronal transfections were performed using Lipofectamine LTX according to the manufacturer’s instructions (Invitrogen, Carlsbad, CA, USA) on 12-14 days *in vitro* (DIV) after plating. Briefly, DNA (0.5 μg/well) was mixed with 0.5 μl PLUS reagent in 50 μl Neurobasal A medium, then mixed with 1.0 μl Lipofectamine LTX in 50 μl NB-A medium, incubated for 20 min and then added to the neurons in NB-A at 37 °C in 5% CO_2_ for 1 h. Neurons were then rinsed with NB-A and incubated in the original medium at 37 °C in 5% CO_2_ for 4-5 days. For co-transfection, neurons were transfected with 1.0 μg of DNA consisting of two plasmids (0.50 μg each).

For HeLa cell culture or HEK293T cell culture, DMEM supplemented with 10% FBS were used. Cell transfections were performed using Lipofectamine2000 according to the manufacturer’s instructions (Invitrogen) after plating.

For inhibitor treatment, HeLa cells or primary neurons cultured on coverslips were pre-incubated with MK801 (10 μM, Sigma-Aldrich), Latruculin A (100 nM, Sigma-Aldrich), NSC 23766 trihydrochloride (100 μM, Abcam), CK-666 (100 μM, Sigma-Aldrich), W-7 (20 μM, TOCRIS), KN-62 (4 μM, TOCRIS), BAPTA-AM (10 μM, Sigma-Aldrich) for 30 min. Neurons were pre-incubated with ML141 (15 μM, Sigma-Aldrich) or SMIFH2 (30 μM, Millipore) for 2 h or with CT04 (2 μg/ml, Cytoskeleton) for 3 h. For tetanus toxin treatment (10 nM, Sigma-Aldrich), neurons were pre-incubated for 10 min. These drugs were maintained during glycine or ionomycin application.

For ionomycin treatment of HeLa cells, cells were pre-incubated with modified Krebs-Ringer Hepes buffer (containing 120 mM NaCl, 4.8 mM KCl, 1.2 mM KH_2_PO_4_, 1.2 mM MgSO_4_, 1.3 mM CaCl_2_, 5.5 mM glucose, 25 mM HEPES, pH 7.4, at 37 °C) for 30 min then treated with ionomycin (2 μM, Beyotime Biotechnology) in KRH buffer for 20 min.

### Chemically-induced LTP (cLTP)

Chemical induction of LTP was performed as previously described (Fortin et al., 2010; Park et al., 2006). Briefly, neurons were treated with glycine (200 μM) in Mg^2+^-free extracellular iso-osmotic solution (mM: 125 NaCl, 2.5 KCl, 2 CaCl_2_, 5 HEPES, 33 glucose, 0.2 glycine, 0.02 bicuculline, and 0.003 strychnine, pH 7.4) (Yang et al., 2018). For experiments performed in the absence of extracellular Ca^2+^, 10 μM BAPTA-AM was substituted for 2 mM CaCl_2_ in the Mg^2+^-free extracellular solution.

For experiment to determine the role of membrane tension in spine expansion, neurons were pretreated with the hypo-osmotic solution (mM: 80 NaCl, 2.5 KCl, 2 CaCl_2_, 5 HEPES, 33 glucose, pH 7.4, OSM 210) or hyper-osmotic solution (mM: 125 NaCl, 2.5 KCl, 2 CaCl_2_, 5 HEPES, 33 glucose, 250 mM sucrose, pH 7.4, OSM 600) for 20 min and chemically induced LTP in the same solution.

### Immunostaining, image acquisition and analysis

Neurons were fixed in 4% PFA/4% sucrose in PBS at RT for 15 min. After blocking with 1% BSA in PBS containing 0.4% Triton X-100 for 40 min at RT, neurons were incubated with primary antibodies for 1 h at RT or overnight at 4°C, and appropriate secondary antibodies conjugated with Alexa Fluor 488, Alexa Fluor 555, or Alexa Fluor 647 were applied for detection.

Confocal images were collected using the Spectral Imaging Confocal Microscope Digital Eclipse C1Si (Nikon, Tokyo, Japan) with a 100× Plan Apochromat VC (NA 1.40) oil objective. Images were z projections of images taken at 0.2 μm step intervals. The number of planes, typically 4-6, was chosen to encompass the entire dendrite from top to bottom.

The procedure for morphometric analysis of dendritic spines was described previously (Yang et al., 2015) (Yang et al., 2018). DsRed was used as a cell-fill. The final reconstructed spines were obtained using a maximum-intensity projection strategy provided by NIS-Elements AR software (Nikon). DsRed-labeled spines were outlined manually. Dendritic segments 40-120 μm from the neuronal cell body were selected for analysis. All morphological experiments were repeated at least three times with an n ≥ 11 for individual experiments.

### Super-resolution live cell imaging and data analysis

The GI-SIM live imaging experiments were performed as described by Guo et al (Guo et al., 2018). Briefly, mouse hippocampal neurons cultured on 25-mm coverslips were transfected with constructs expressing membrane-bound GFP (mGFP) and LifeAct-mCherry on DIV12 and imaged on DIV16 in Mg^2+^-free extracellular solution (Yang et al., 2018). Time-lapse images were obtained with acquisition time of 110 ms for each channel at 5 s intervals. To quantify the area of each spine head and enrichment of F-actin in spines, we measured the fluorescence mean intensity of LifeAct-mCherry within the spines and normalized each measurement by the fluorescence signal along the adjacent dendritic shaft with the NIH ImageJ software. The mGFP-labeled dendrites or spines were outlined manually.

For analysis of the spatiotemporal relationship between actin polymerization and membrane expansion in dendritic spines, spines were segmented from raw GI-SIM images using Otsu’s method (Otsu, 1979). The area of each spine and the mean fluorescence signal of F-actin inside each spine were then quantified using NIH ImageJ. To facilitate the visualization of instantaneous spine growth and localized actin polymerization, differential images of both membrane (mGFP) and actin (LifeAct-mCherry) channels were calculated by subtracting the image at time point *t* from that at *t+1* through the entire time lapse movie (1 min duration before and 3 min duration post glycine application with 5 s intervals) using Matlab (R2018a, Mathworks). To highlight the regions with spine growth or increased F-actin signals, only pixels with a positive difference were displayed in the final differential images. We then measured the overlap between differential images in the membrane channel and those in the actin channel at individual time points, and determined the extent of overlap by calculating the ratio of overlapped to total changes in the membrane channel using the JACoP plug-in of the ImageJ software. Then we obtained fluctuations in the extent of overlap at different time points before or post glycine treatment to evaluate the randomness of overlap between membrane expansion and actin polymerization.

### Immunohistochemical analyses

Mice were anesthetized with 1% sodium pentobarbital and transcardially perfused with normal saline followed by 4% paraformaldehyde (PFA) in 0.01 M phosphate-buffered saline (PBS). Mouse brain was dissected out and post-fixed with 4% PFA/PBS for 4 h at 4 °C. Fixed brain was incubated with 20% sucrose overnight and then 30% sucrose overnight. The brain was embedded in OCT and stored at −80°C until usage. Thirty-micron cyrosections were made using cryostat and collected.

For immunostaining of brain sections, floating 30 μm-thick slices were rinsed with PBS and permeabilized in 0.4% Triton X-100 in 0.01M PBS for 30 min. Cyrosections were blocked with 1% BSA in PBS containing 0.4% Triton X-100 for 1 h at RT, then incubated with primary antibodies overnight at 4 °C. Secondary antibodies conjugated with Alexa Fluor 555 were used for detection. Sections were then incubated with DAPI (Roche, Grenzach-Wyhlen, Germany) for nuclear staining for 5 min at RT. Following rinsing, cyrosections were mounted on gelatin-coated slides and covered with coverslip with mounting medium. Confocal images were collected using the Spectral Imaging Confocal Microscope Digital Eclipse C1Si (Nikon, Tokyo, Japan) with a 10×Plan Apochromat DIC N1 0.45 objective or 40×Plan Fluo (NA 1.30) oil objective (Yang et al., 2018).

### Protein expression and purification

His-EEN1, His-calmodulin, His-p140Cap fragment (aa 351-1051) or GST-Calmodulin was expressed in *E. coli* BL21 (DE3). Cells were grown at 37 °C in LB (g/L: tryptone 10, yeast extract 5, NaCl 10) supplemented with ampicillin or kanamycin. Cells were induced at OD_600_ of ~0.6 with 0.4 mM isopropyl β-D-1-thiogalactopyranoside (IPTG) for 4 h at 30 °C or 16 h at 16 °C. Cells were harvested and stored at −80 °C until purification.

For His-tagged proteins, cells were resuspended in lysis buffer (50 mM NaH_2_PO_4_, 300 mM NaCl, 15 mM imidazole, pH 8.0) supplemented with 1% Triton X-100 and 0.1 mM PMSF. Protein were purified with Ni-NTA resin according to the manufacturer’s instructions (R90115, Invitrogen). For GST-tagged proteins, cells were resuspended in PBS supplemented with 0.2% Triton X-100 and 0.1 mM PMSF. Protein were purified with Glutathione Sepharose 4B (GE17-0756-01, Sigma-Aldrich, St. Louis, MO, USA) according to the manufacturer’s instructions.

For liposome sedimentation assay, purified His-calmodulin was dialyzed in a Spectra/Por™ 4 RC Dialysis Membrane Tubing (08-667D, Thermo Fisher Scientific, Waltham, MA, USA) against 1,000 volumes of PBS supplemented with 0.5 mM EGTA at 4 °C for 4 h and replaced with fresh 1,000 volumes of PBS twice.

### Isothermal titration calorimetry (ITC)

ITC was performed on a MicroCal PEAQ-ITC ((Malvern Panalytical, U.S.A) calorimeter. Calmodulin and EEN1 were purified on a Superdex-200 16/600 column (GE Healthcare, U.S.A) in solution buffer containing 20 mM Tris-Cl at pH 7.0, 150 mM NaCl. Solution buffer containing 1 mM calcium was injected into the calorimeter cell fullfilled with protein1solution, cell temperature set to 25°C. Each analysis involved 20 injections of 4 s duration (2 μL per one injection), 120 s spacing, stir with 750 rpm, 5 μcal/s reference power and high gain feedback mode. Data were processed by Origin software to obtain thermodynamic profiles.

### GST-pull down, co-immunoprecipitation (IP) and immunoisolation

For GST-pull down assays, 5 μg of GST-tagged protein conjugated with glutathione-Sepharose beads was incubated with 1 μg of His-tagged protein in 0.01 M PBS supplemented with 1% NP-40 at 4°C for 1 h. Beads then was washed five times with PBS supplemented with 0.3% Triton-X 100 and was boiled in SDS sample buffer.

For co-IP experiments, HEK293T cells, cultured neurons or mouse brain were lysed with lysis buffer 1 (0.05% [vol/vol] NP-40, 15 mM Tris-HCl, pH 7.4, 50 mM NaCl) supplemented with protease inhibitors for IP of endogenous proteins (endo-IP), or with lysis buffer 2 (0.1% [vol/vol] NP-40, 50 mM Tris-HCl, pH 7.4, 150 mM NaCl) supplemented with protease inhibitors for Flag-IP. Lysates were then centrifuged at 16,000 × g for 15 min at 4°C. For Flag IP, cell lysates were incubated with anti-Flag Affinity Gel (Sigma-Aldrich) at 4°C for 4 h on a roller mixer. For endo-IP, antibody (1 μg) was added to the cell lysates and incubated at 4◻ for 2 h on a roller mixer, followed by incubation with Protein G agarose (Santa Cruz) pre-equilibrated in lysis buffer overnight at 4°C. Immunoprecipitates were washed four times in lysis buffer and boiled in SDS sample buffer, then subjected to SDS-PAGE s and immunoblotting.

For immunoisolation of membrane proteins, cultured neurons were homogenized with lysis buffer (mM: Tris-HCl 20, HEPES pH 7.4 10, NaCl 150, sucrose 250) supplemented with protease inhibitors and centrifuged at 800 × g for 15 min. The supernatants were collected and subjected to high-speed centrifugation at 100,000 × g for 1h (TLS-55 rotor, OptimaTMMAX Ultracentrifuge; Beckman Coulter, Germany). The supernatants (S100) and pellets (p100, the membrane fraction) resuspended in lysis buffer were subjected to immunoisolation with Dynabeads Protein G (Invitrogen, Carlsbad, CA, USA) coupled with 2 μg of mouse anti-calmodulin antibody. Bound proteins were eluted by boiling in 2× SDS gel loading buffer and subjected to SDS-PAGE and immunoblotting.

### Subcellular fractionation

Cultured neurons or mouse hippocampi were homogenized to isolate the membrane and cytoskeleton fractions with Subcellular Protein Fractionation Kit for Cultured Cells (Thermo, 77840) or for tissues (Thermo, 87790) according to the manufacturer’s instructions. Proteins in different fractions were subjected to SDS-PAGE and immunoblotting.

### Liposome co-sedimentation assay

Brain extract from bovine brain (Sigma-Aldrich, B1502) were dissolved in chloroform and dried under vacuum for 30 min. The solvent-free lipid films were rehydrated with liposome buffer (150◻mM NaCl, 20◻mM Tris-HCl pH 7.4, 1 mM DTT) and subjected to 7 cycles of flash freezing in liquid nitrogen and thawing in a 37°C bath. Liposomes were then extruded 21 times through a polycarbonate membrane with a 50◻nm pore size (Mini-Extruder, Avanti Polar Lipids). Extruded liposomes were centrifuged at 18,000 g for 5 min to remove insoluble material and stored at 4°C. Liposomes (0.5 mg/ml) were then incubated with freshly purified recombinant proteins (1 μM His-EEN1 or 1 μM His-EEN1 and 2 μM His-calmodulin), in 100◻μl liposome buffer for 10 min at 30°C before sedimentation at 140,000g for 30 min at 4°C. The supernatant (unbound) and pellet (bound) were subjected to SDS-PAGE. Ratios of binding to liposomes were determined using NIH ImageJ software.

### Three-dimensional Structured Illumination Microscopy (3D-SIM) imaging and image analysis

3D-SIM images were acquired as previously described (Niu et al., 2013) on the DeltaVision OMX V4 imaging system (Applied Precision Inc, USA) with a 100 × 1.4 oil objective (Olympus UPlanSApo), solid state multimode lasers (488, 593 and 642 nm) and electron-multiplying CCD (charge-coupled device) cameras (Evolve 512×512, Photometrics, USA). Serial Z-stack sectioning was done at 125 nm intervals. The microscope is routinely calibrated with 100 nm fluorescent spheres to calculate both the lateral and axial limits of image resolution. SIM image stacks were reconstructed using softWoRx 5.0 (Applied Precision) with the following settings: pixel size 39.5 nm; channel-specific optical transfer functions; Wiener filter 0.001000; discard Negative Intensities background; drift correction with respect to first angle; custom K0 guess angles for camera positions. Pixel registration was corrected to be less than 1 pixel for all channels using 100 nm Tetraspeck beads. For clarity of display, small linear changes to brightness and contrast were performed on three-dimensional reconstructions.

### Electrophysiology

*EEN1^fl/fl^* mice within 24 h after birth were co-injected with high-titer AAV stock carrying pAOV-CAMKII-mCherry-2A-Cre (AAV-mCherry-2A-Cre) and pAAV-CaMKIIa-EGFP-2A-MCS-3FLAG-EEN1 or mutants of EEN1 (about 1 ~ 5 × 10^13^ IU/ml). Newborns were anesthetized on ice for 5 minutes and then mounted in a custom ceramic mold to make the head level in the X – and Y – axes. Lambda was set as (X, Y) = (0, 0). Zero point of Z – axis was the position at which the injecting needle penetrated the skin. About 10 nl viral solution was injected at each of the seven sites ((X, Y, Z) = (1.2, 1.2, 1.4/ 1.0/ 0.6) and (1.5, 1.0, 1.7/ 1.3/ 0.9/ 0.5)) targeting the hippocampus at each cerebral hemisphere with microsyringe (Sutter Instrument) and a beveled glass injection pipette. Injected pups were returned to home cage and used for recording two to three weeks afterward. Transverse 350 μm hippocampal slices were cut from viral injected *EEN1^fl/fl^* mice on a Leica vibratome (VT1000 S) in high sucrose cutting solution containing (in mM): KCl 2.5, NaH_2_PO_4_ 1.25, NaHCO_3_ 25, CaCl_2_ 0.50, MgSO_4_ 7, sucrose 210, glucose 10, Na-ascorbic acid 1.3. Freshly cut slices were placed in an incubating chamber containing ACSF, and recovered at 32°C for about 20 min followed by 60 min at room temperature before recording. The slices were perfused with ACSF containing GABAA receptor antagonists PTX (100 μM) / Bic (10 μM) and saturated with 95% O_2_/5% CO_2_ in whole-cell LTP experiments. CA1 pyramidal cells were voltage-clamped at −70 mV and AMPAR EPSCs were evoked by stimulation at SC with concentric electrode (FHC CBBRC75). LTP was induced by stimulating SC axons at 2 Hz for 90 s while clamping the cell at 0 mV, after recording a stable 3- to 5-min baseline, but no more than 6 min after breaking into the cell (Diaz-Alonso et al., 2017; Granger et al., 2013). To minimize run-up of baseline responses during LTP, cells were held cell-attached for about 1 to 2 min before breaking into the cell.

### Stereotaxic injection and Behavioral tests

Nine-week-old sexually naive male and female mice were anesthetized and stereotactically injected with viral particles in the hippocampal CA1 region as described (Yang et al., 2018). The virus-injected mice were tested for behavior two weeks later. Morris water maze and fear conditioning tests were performed as previously described (Yang et al., 2018). We observed no sex-related difference in behaviors and the results were pooled together.

### Statistical analysis

All data were presented as the mean ± SEM. GraphPad Prism 5 (GraphPad Software, LaJolla, CA) was used for statistical analysis. For two-sample comparisons vs. controls, unpaired Student’s t-test was used except where noted. One-way analysis of variance with a Dunnett’s multiple-comparison or Newman-Keuls multiple comparison *hoc* test was used to evaluate statistical significance of three or more groups of samples. A *p* value of less than 0.05 was considered statistically significant.

## Additional files

Supplementary files:

(A) Figure supplements

**Figure 3 – figure supplement 1.** Endophilin A1 does not bind Ca^2+^.

**Figure 4 – figure supplement 1.** I154 and L158 are required for endophilin A1 binding to calmodulin.

**Figure 5 – figure supplement 1.** Calmodulin-dependent increase in the number of endophilin A1 nanodomains correlates with spine size during the initial phase of sLTP.

**Figure 6 – figure supplement 1.** In response to Ca^2+^/calmodulin, endophilin A1 localizes to the plasma membrane and recruits Arp2/3 via p140Cap.

(B) Supplementary Videos

**Video 1.** Morphological changes and actin dynamics in *EEN1* WT dendrite before and during glycine-induced sLTP.

Dendrites of DIV16 *EEN1* WT mouse hippocampal neurons co-expressing mGFP (green) and the F-actin probe LifeAct-mCherry (red) were imaged every 5 s by GI-SIM for 12 frames before and 48 frames after glycine treatment. Video plays at 5 frames/s. Scale bars, 5 μm (left) and 1 μm (right).

**Video 2.** Morphological changes and actin dynamics in *EEN1* KO dendrite before and during glycine-induced sLTP.

Dendrites of DIV16 *EEN1* KO mouse hippocampal neurons co-expressing mGFP (green) and the F-actin probe LifeAct-mCherry (red) were imaged every 5 s by GI-SIM for 12 frames before and 48 frames after glycine treatment. Video plays at 5 frames/s. Scale bars, 5 μm (left) and 1 μm (right).

**Video 3.** Spatiotemporal relationship of spine growth and actin polymerization in *EEN1* WT spines before and during glycine-induced LTP.

Increases in spine head area and F-actin signal intensity are color coded green and red, respectively. Video plays at 5 frames/s. Scale bars, 5 μm (left) and 1 μm (right).

**Video 4.** Spatiotemporal relationship of spine growth and actin polymerization in *EEN1* KO spines before and during glycine-induced LTP.

Increases in spine head area and F-actin signal intensity are color coded green and red, respectively. Video plays at 5 frames/s. Scale bars, 5 μm (left) and 1 μm (right).

