## Supplemental figures for "Endophilin A1 promotes Actin Polymerization in response to Ca^2+^/calmodulin to Initiate Structural Plasticity of Dendritic Spines"

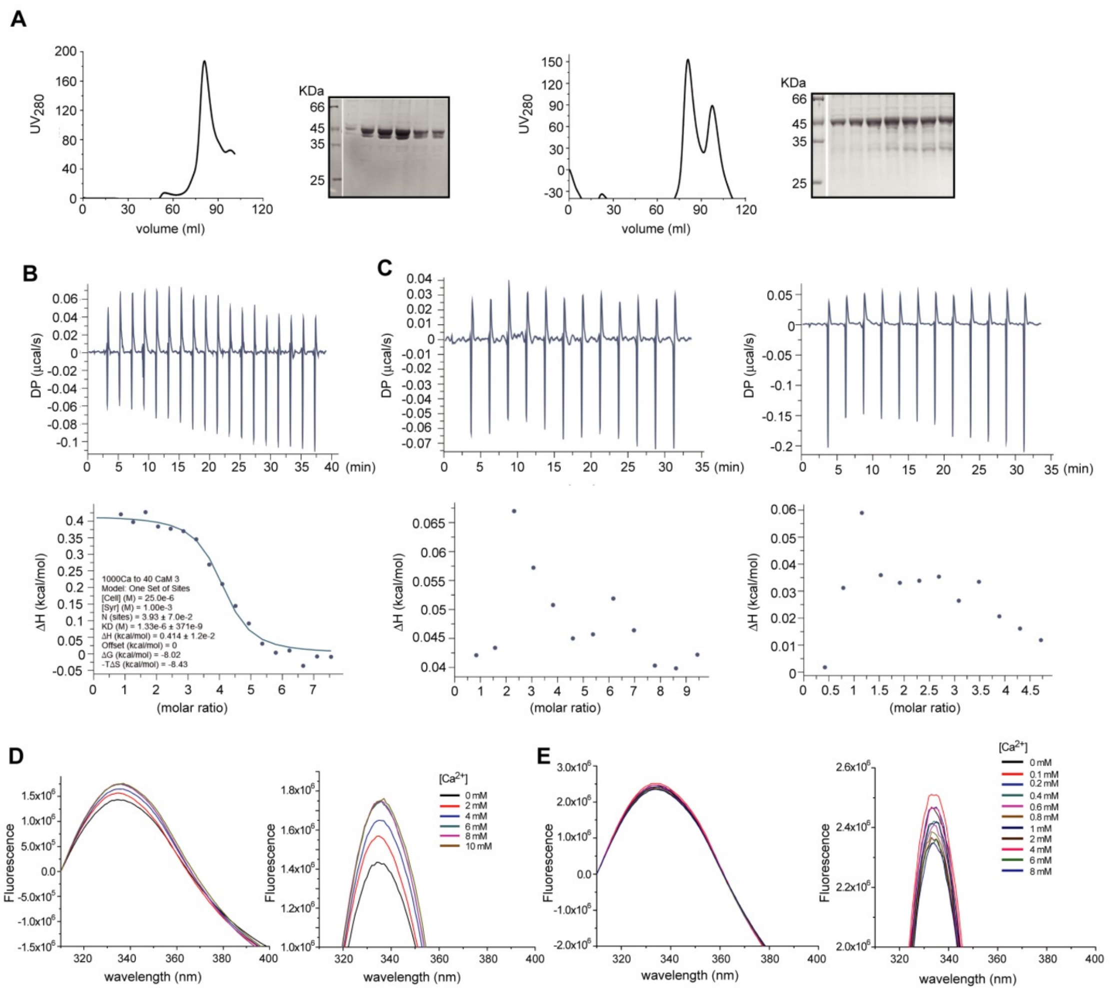


**Figure 3 —figure supplement 1.** Endophilin A1 does not bind $Ca^{2+}$.

(**A**) Purification of recombinant full-length EEN1 expressed in *E.coli*. Affinity-purified His-tagged EEN1 was supplied with 5 mM EDTA and loaded into Superdex 200 16/300 size exclusion chromatography with buffer (20 mM Tris-Cl pH 7.0, 150 mM NaCl). The peak fractions for EEN1 were collected and analyzed by SDS-PAGE. EEN1 displays as monomer and presents negligible degradation without affecting its function. (**B**) ITC of 1 mM $CaCl_2$ and 25 µM CaM. Measured disassociation constant is 1.33 ± 0.37 µM with four binding sites. The enthalpy change (ΔH) of the reaction is 0.414 ± 0.012 kcal·$mol^{-1}$, which indicates as an endothermic reaction. The entropy of the reaction system is increased, suggesting conformational change of CaM. (**C**) ITC of 1 mM $CaCl_2$, 40 µM EEN1 (left panels) and 3 mM $CaCl_2$, 100 µM EEN1 (right panels). No significant binding was detected. (**D, E**) Tryptophan fluorescence of Synaptotagmin-1 C2B (served as positive control) and EEN1 in the presence of $CaCl_2$. Shown are tryptophan fluorescence spectra (310-400 nm) of Synaptagmin-1 C2B (**D**) and EEN1 (**E**) respectively with an excitation wavelength of 280 nm.

Right panels are enlarged views of the emission peaks in the left panels. The tryptophan fluorescence of Synaptagmin-1 C2B increases with the addition of $CaCl_2$ and saturates at 6 mM final concentration. In contrast, the tryptophan fluorescence of EEN1 fluctuates irregularly with the addition of $CaCl_2$.

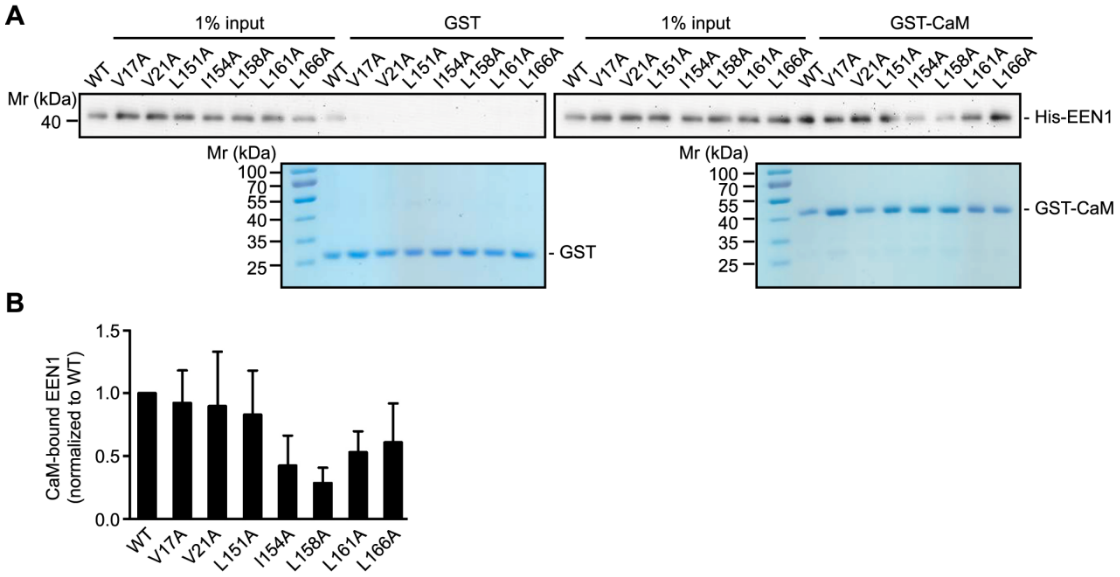


**Figure 4**一**figure supplement 1.** I154 and L158 are required for endophilin A1 binding to calmodulin.

(**A**) Binding of endophilin A1 WT and mutants to GST-CaM in pull-down assay. Bound proteins were analyzed by SDS-PAGE and immunoblotting with anti-His antibodies. (**B**) Quantification of mutants binding to GST-CaM in (**A**). Data represent mean ± SEM, N = 5 independent experiments.

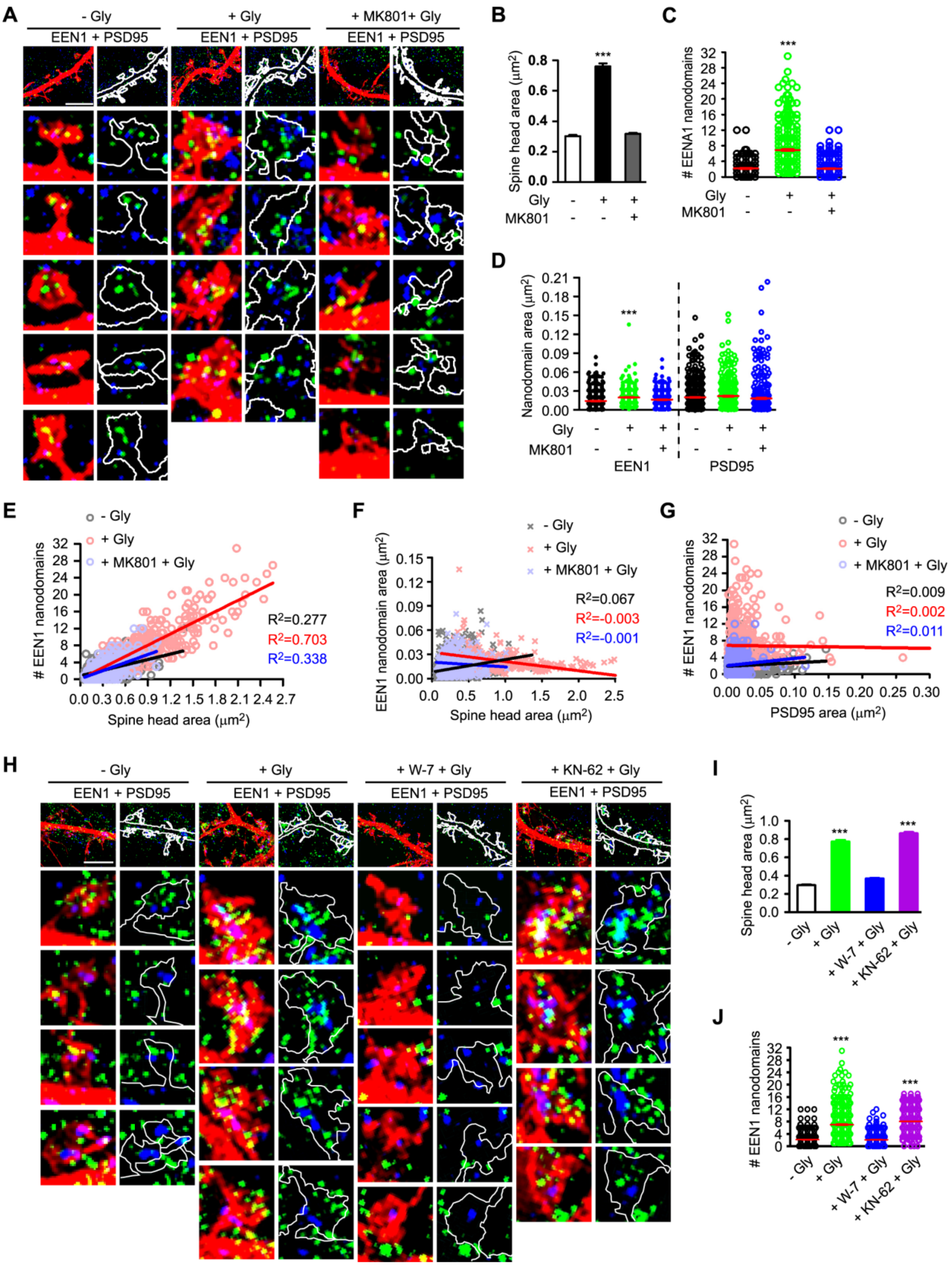


**Figure 5 —figure supplement 1.** Calmodulin-dependent increase in the number of endophilin A1 nanodomains correlates with spine size during the initial Phase of sLTP.

(**A**) Cultured hippocampal neurons transfected with pLL3.7-DsRed on DIV12 were pre-treated with DMSO or MK801 and chemically induced LTP on DIV16. Neurons were fixed 5 min after glycine application, immunostained with antibodies to endophilin A1 (green) and PSD95 (blue) and imaged by 3D-structured illumination microscopy (SIM). Shown are representative images. Spines were outlined manually. Bar = 4 µm. (**B**) Quantification of spine size in (**A**). N ≥ 13 neurons, n ≥ 502 spines per group. (**C**) Quantification of the number of endophilin A1 nanodomains in spines in (**A**). N ≥ 13 neurons, n ≥ 502 spines per group. (**D**) Quantification of the area of endophilin A1 or PSD95 nanodomains in spines in (**A**). N ≥ 13 neurons, n ≥ 502 spines per group. (**E, F**) Scatterplot of the number or area of endophilin A1 nanodomains versus the size of spine head with linear fits. (**G**) Scatterplot of the number of endophilin A1 nanodomains versus the area of PSD95 nanodomains in spines. (**H**) Effect of W-7 or KN-62 on endophilin A1 nanodomains in spines. Neurons were pre-treated with DMSO, W-7 or KN-62 and chemically induced LTP on DIV16. Neurons were fixed 5 min after glycine application, immunostained and imaged by 3D-SIM. Shown are representative images. Bar = 4 µm. (**I**) Quantification of spine size in (**H**). N ≥ 14 neurons, n ≥ 540 spines per group. (**J**) Quantification of the number of endophilin A1 nanodomains in spines in (**H**). N ≥ 14 neurons, n ≥ 540 spines per group. Data represent mean ± SEM. *** $p < 0.001$ when compared with - Gly.

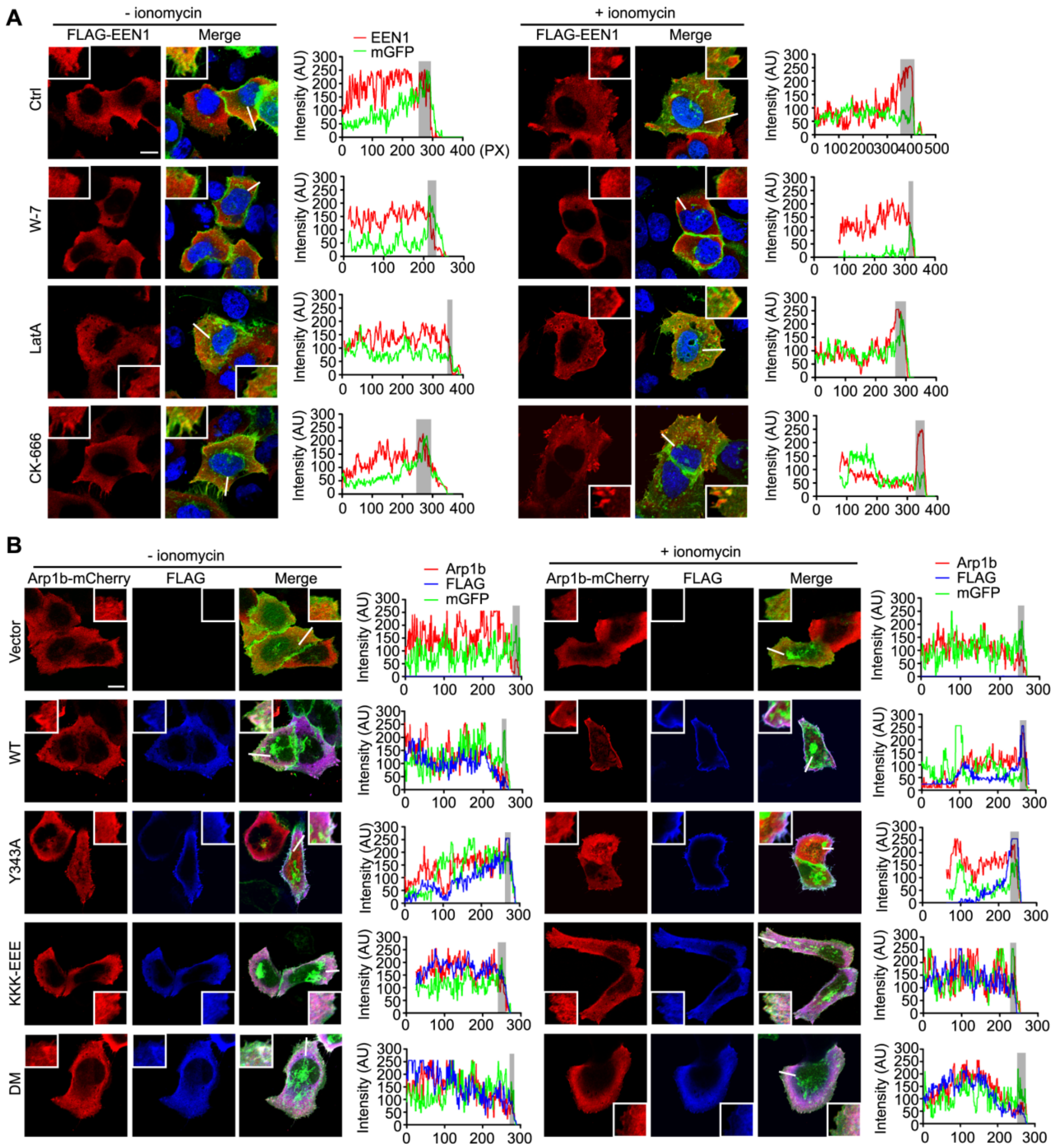


**Figure 6 —figure supplement 1.** In response to $Ca^{2+}$/calmodulin, endophilin A1 localizes to the plasma membrane and recruits Arp2/3 via p140Cap.

(**A**) HeLa cells were co-transfected with constructs expressing mGFP, myc-p140Cap and Flag-EEN1 for 24 h, then pretreated with DMSO, W-7, LatA or CK-666 for 30 min followed by ionomycin treatment. Cells were fixed 20 min after ionomycin treatment and imaged by confocal microscopy. Left panels are representative images. Magnified images are shown in white boxes. Right panels are linescan plots of the fluorescence intensity along the white lines indicated in the images. Grey shaded areas indicate mGFP-labeled plasma membrane. AU: arbitrary unit. PX: pixels. Bar = 10 μm. (**B**) HeLa cells were co-transfected with constructs expressing mGFP, myc-p140Cap, Arp1b-mCherry and Flag-EEN1 WT, Y343A, KKK-EEE, or DM mutant for 24 h, then treated with ionomycin. Cells were fixed 20 min after ionomycin treatment and imaged by confocal microscopy. Left panels are representative images. Magnified images are shown in white boxes. Right panels are linescan plots of the fluorescence intensity along the white lines indicated in the images. Grey shaded areas indicate mGFP-labeled plasma membrane. Bar = 10 μm.
